# Oxidized EGCG redirects α-synuclein aggregation through shear-selected interactions

**DOI:** 10.64898/2026.10.25.761737

**Authors:** Cho Ming Yuen, Reiji Yamaguchi, Keita Uchida, Yosuke Shimada, Yusuke Shuto, Shunta Sakamoto, Daichi Morimoto, Hajime Ono, Masatomo So, Ayako Furukawa, Kenji Sugase

## Abstract

α-Synuclein (αSyn) aggregation is central to Parkinson’s disease pathology, and the green-tea polyphenol epigallocatechin gallate (EGCG) has been studied as an inhibitor of fibril formation. However, EGCG autoxidation and analyses under static, non-fibrillating conditions have obscured whether activity arises from EGCG itself or oxidized species and which interactions persist during αSyn assembly. Here we combine HPLC, LC-MS, TEM and solution NMR to compare unoxidized and oxidized EGCG (oxEGCG), while Rheo-NMR monitors αSyn during shear-promoted assembly. Unoxidized EGCG showed no detectable binding under static, reducing conditions, whereas oxEGCG bound αSyn, inhibited fibrillation and remodeled preformed fibrils. Under shear flow, many αSyn chemical shifts approached their free-state values, whereas interactions near the NAC–C-terminal boundary were preferentially retained, coinciding with formation of oligomers and short, thick fibrils. These findings identify oxidation state and flow as determinants of EGCG action on αSyn and show that Rheo-NMR distinguishes interactions that persist during shear-promoted assembly.

---

Parkinson’s disease is a progressive neurodegenerative disorder characterized by the accumulation and propagation of misfolded α-synuclein (αSyn). Lewy bodies, the pathological hallmark of the disease, contain αSyn amyloid fibrils as a major component^1,2^, whereas soluble oligomeric intermediates formed during aggregation have been implicated in cytotoxicity and neuronal dysfunction^3,4^. Because both oligomeric intermediates and fibrillar aggregates are associated with neurotoxicity and pathological propagation, understanding the factors that direct αSyn into distinct assembly pathways may inform strategies for mitigating αSyn-associated neurotoxicity and disease progression^5,6^.

Among potential modulators of αSyn aggregation, tea has attracted attention because epidemiological studies have associated its consumption with a reduced risk of Parkinson’s disease^7,8^. The beneficial effects of green tea have often been attributed to epigallocatechin gallate (EGCG), a catechin polyphenol abundant in tea leaves. In the context of αSyn, EGCG has been proposed to bind monomeric or aggregated species and redirect aggregation toward off-pathway oligomeric assemblies rather than mature amyloid fibrils^9^. EGCG has also been reported to remodel preformed amyloid fibrils into non-fibrillar aggregates, suggesting that it can affect both fibril formation and the stability of existing amyloid assemblies^10^. Similar anti-amyloid effects have been reported for other amyloidogenic proteins, including amyloid-β, islet amyloid polypeptide and lysozyme^9,11,12^. These diverse effects suggest that EGCG may alter amyloid assembly pathways rather than simply block fibril formation. However, the molecular basis of these effects remains unclear in two important respects.

First, EGCG readily undergoes autoxidation under aerobic assay conditions, producing a heterogeneous and time-dependent mixture of oxidation products during the course of amyloid assays^13,14^. Consequently, binding, fibrillation inhibition and fibril remodeling attributed to EGCG may arise from the parent catechin, oxidized species generated during the experiment, or both. Resolving this distinction is essential for understanding how EGCG modulates αSyn aggregation.

Second, it remains unclear which EGCG–αSyn interactions are relevant while αSyn is actively assembling. Previous NMR studies have characterized EGCG–αSyn interactions at residue-level resolution under static conditions^15,16^. However, at neutral pH, αSyn does not readily form amyloid fibrils under conventional static NMR conditions^17^. Static measurements therefore cannot distinguish interactions that persist during aggregation from those that are lost once assembly begins. Resolving this question requires residue-level monitoring of molecular interactions under conditions in which aggregation is actively proceeding.

Here, we addressed these two questions by defining the oxidation dependence of EGCG activity using HPLC, LC-MS, TEM and solution NMR, and then using Rheo-NMR to identify oxidized EGCG (oxEGCG)–αSyn interactions that persist during shear-promoted assembly^18–21^. We show that detectable αSyn binding, fibrillation inhibition and fibril remodeling arise primarily from oxidized EGCG species, whereas unoxidized EGCG shows no detectable interaction under static, reducing conditions. Under shear flow, many oxEGCG-dependent perturbations are partially reversed during assembly, whereas interactions near the NAC–C-terminal boundary are preferentially retained, coinciding with the formation of oligomers and short, thick fibrils.

## Results

### Oxidized EGCG inhibits αSyn fibrillation and remodels preformed fibrils

Incubation of an EGCG solution at 25 °C for up to 48 h resulted in gradual darkening and the appearance of multiple chromatographic peaks (Fig. 1a), indicating the formation of a heterogeneous mixture of oxidation products (hereafter oxEGCG). These components showed complex time-dependent changes, with some peaks increasing in intensity whereas others decreased or disappeared, reflecting dynamic EGCG autoxidation. Known EGCG oxidation products were detected, including gallic acid and species consistent with theasinensins A and D (Extended Data Fig. 1a–c and Supplementary Section 1). A small gallic acid peak was also detected at 0 h (Fig. 1a), suggesting that trace amounts were either initially present or formed during sample preparation. Several peaks remained unidentified, and the instability of the oxidation products complicated their isolation and structural identification. Given this heterogeneity and instability, we used unfractionated oxidized EGCG preparations in subsequent experiments rather than isolated oxidation products.

**Fig. 1.**
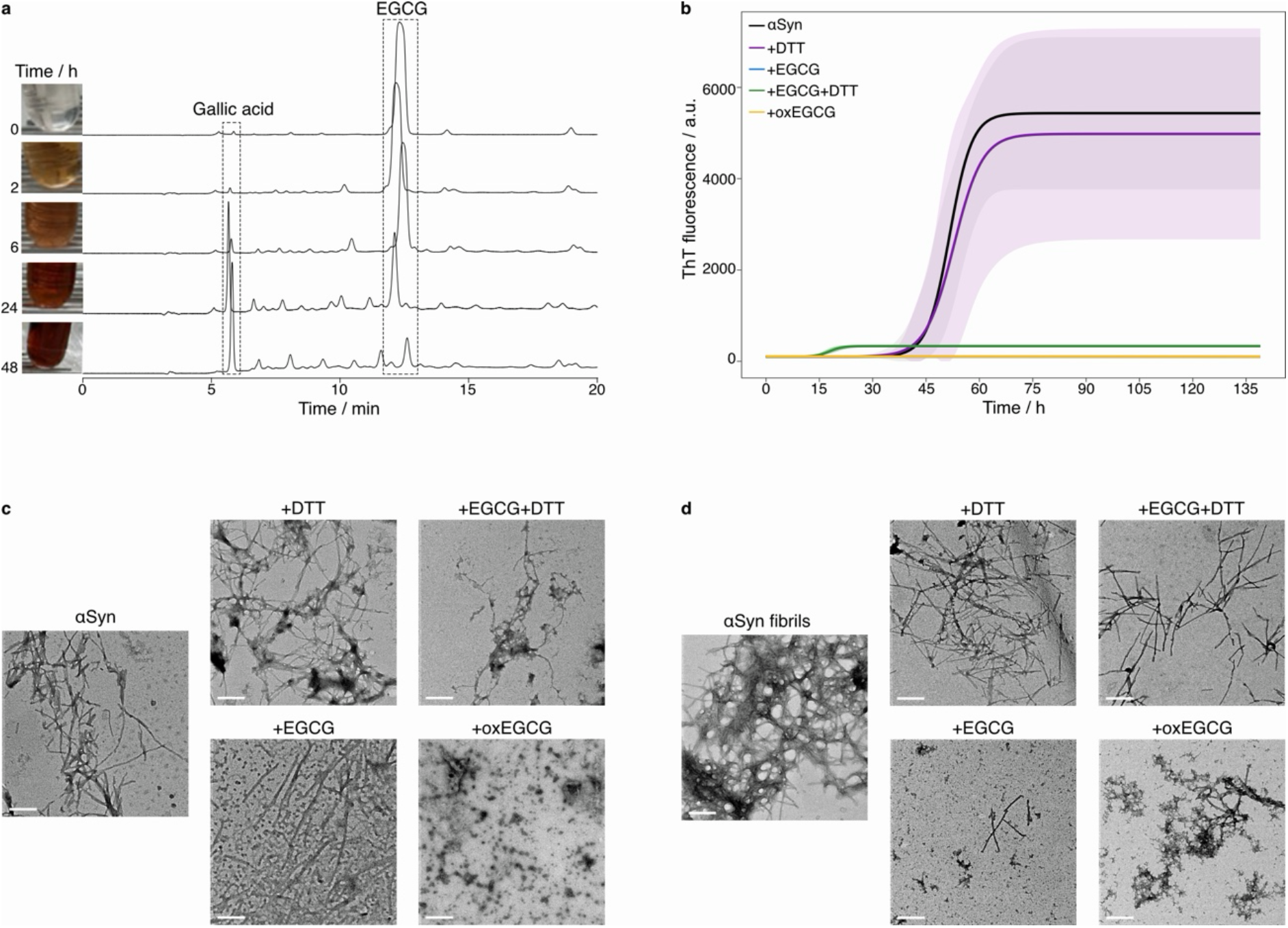
| Characterization of oxEGCG and its effects on αSyn fibrillation and fibril remodeling. **a**, Time-dependent color changes of the EGCG solution and corresponding HPLC chromatograms. **b**, αSyn fibrillation monitored by ThT fluorescence in buffer alone or buffer supplemented with oxEGCG, EGCG, EGCG + DTT, or DTT alone. Lines show representative fitted curves; fitting and model selection are described in Supplementary Section 2. Shaded areas indicate ± s.d. The numbers of fluorescence traces assigned to each model are provided in Extended Data Table 1. **c**,**d**, Representative TEM images of αSyn aggregates after the ThT assay (**c**) and preformed fibrils after incubation (**d**) in buffer alone or buffer supplemented with oxEGCG, EGCG, EGCG + DTT, or DTT alone. Scale bar, 300 nm.

We then monitored αSyn fibril formation by thioflavin T (ThT) fluorescence in buffer alone or buffer supplemented with oxEGCG (oxidized for 72 h), EGCG, EGCG + DTT, or DTT alone (Fig. 1b). In the buffer control, ThT fluorescence increased in a sigmoidal manner, consistent with αSyn amyloid fibril formation (Supplementary Section 2, Extended Data Fig. 2 and Extended Data Table 1). In contrast, oxEGCG, EGCG, and EGCG + DTT all suppressed the increase in ThT fluorescence. Among these conditions, oxEGCG and EGCG yielded the lowest final fluorescence intensities and showed nearly overlapping profiles close to the fluorescence baseline (∼100 a.u.), whereas EGCG + DTT showed a slightly higher fluorescence intensity. HPLC analysis of the EGCG + DTT sample after the ThT assay showed that the EGCG peak was barely detectable (Extended Data Fig. 1d), indicating extensive EGCG oxidation despite the presence of DTT. DTT alone did not substantially alter the ThT profile. These results show that EGCG underwent oxidation during the assay even in the presence of DTT; however, the reduced ThT fluorescence alone does not establish whether αSyn fibril formation was inhibited.

To assess αSyn aggregate morphology independently of ThT fluorescence, we examined the aggregates formed during the ThT assay by transmission electron microscopy (TEM) (Fig. 1c). The buffer control showed typical fibrils, and aggregates formed in the presence of DTT alone were similar to those in the buffer control. In contrast, incubation with oxEGCG produced spherical oligomers rather than fibrils, consistent with previous studies^9,22^. EGCG produced a mixture of fibrillar and oligomeric species. In the presence of EGCG + DTT, fibrils were still observed, whereas the abundance of oligomeric species appeared lower than in the EGCG condition, suggesting that reducing conditions attenuated the effect of EGCG oxidation on αSyn aggregation. Although fibrils were observed in both the EGCG and EGCG + DTT samples, an increase in ThT fluorescence was observed only for EGCG + DTT. This discrepancy was consistent with previous reports that αSyn aggregates formed in the presence of EGCG can exhibit low ThT fluorescence intensity and that EGCG can interfere with ThT detection by competing with ThT for overlapping binding sites on αSyn fibrils^13,23^. Together, the ThT and TEM results show that oxidation enhances the ability of EGCG to redirect αSyn away from typical amyloid fibril formation.

Because oxEGCG altered aggregate morphology during fibril formation, we next examined whether it also altered preformed αSyn fibrils. Preformed fibrils were incubated for 24 h in buffer alone or buffer supplemented with oxEGCG, EGCG, EGCG + DTT, or DTT alone (Fig. 1d). Fibrils were largely retained in the buffer control and in the presence of DTT alone. EGCG appeared to reduce the abundance of fibrillar species, although short fibrils remained. This effect was attenuated by DTT. In contrast, oxEGCG caused marked remodeling of preformed fibrils, leaving non-fibrillar amorphous aggregates. These aggregates were morphologically distinct from the spherical oligomers formed during the fibrillation inhibition assay (Fig. 1c). Collectively, the fibrillation inhibition and fibril remodeling assays indicate that EGCG oxidation modulates both αSyn fibril formation and the remodeling of preformed fibrils.

### Oxidation-dependent interactions of EGCG with αSyn

Because EGCG underwent extensive oxidation during the ThT assay, it remained unclear whether unoxidized EGCG itself interacts with αSyn to inhibit fibrillation. To test this, we acquired ^1^H-^15^N heteronuclear single quantum coherence (HSQC) spectra of αSyn in the absence and presence of EGCG, with DTT added to suppress oxidation. No chemical shift changes were observed initially, but they became detectable after 70 h (Extended Data Fig. 3), suggesting negligible interaction of unoxidized EGCG with αSyn under static, reducing conditions and gradual EGCG oxidation during prolonged incubation. In contrast, oxEGCG titration induced concentration-dependent chemical shift changes, causing most αSyn signals to shift in a similar direction, mainly toward the upper-right (Fig. 2a, left and 2b); the resulting average apparent dissociation constant (*K*_D_) was estimated at 3.2 ± 3.6 mM (Supplementary Figs. 1–3). Because oxEGCG was a heterogeneous mixture and its concentration was defined based on the initial EGCG concentration, the concentration of the active species was unknown and likely lower than the nominal value. The fitted *K*_D_ should therefore be regarded as an apparent value for the mixture rather than the affinity of a defined oxidized species. Analysis of per-residue chemical shift differences (Δ*δ*) showed that oxEGCG produced broad perturbations across the αSyn sequence, with larger effects in the N-terminal region than in the NAC and C-terminal regions (Fig. 2c). Furthermore, several of the largest perturbations were observed for alanine and valine residues (Extended Data Fig. 4a). No macroscopic precipitation was visible in the NMR tube after the oxEGCG titration.

**Fig. 2.**
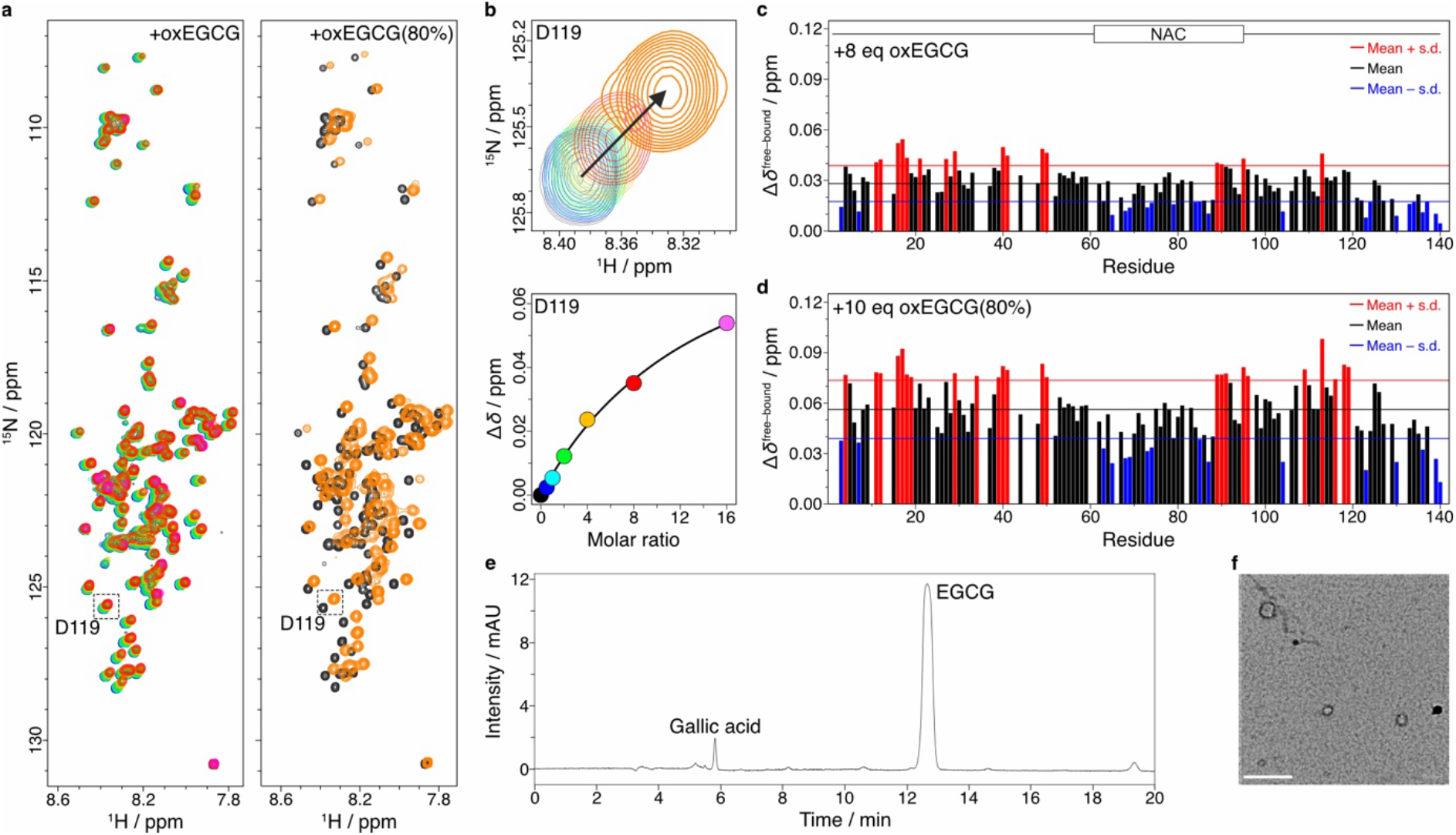
| NMR analysis of oxEGCG–αSyn interactions. **a**, Left, overlay of ^1^H-^15^N HSQC spectra of αSyn in the presence of laboratory-prepared oxEGCG at 0 (black), 0.5 (blue), 1 (cyan), 2 (green), 4 (yellow), 8 (red) and 16 (magenta) equivalents; right, overlay of spectra of αSyn at 0 (black) and 10 (orange) equivalents of the commercial EGCG reagent (oxEGCG(80%)). **b**, Enlarged view of a representative spectral region and the corresponding fit to a 1:1 binding model for D119. **c**, Residue-specific weighted chemical shift differences (Δ*δ*) of αSyn in the presence of 8 equivalents of laboratory-prepared oxEGCG. **d**, Residue-specific Δ*δ*^free–bound^ values of αSyn in the presence of 10 equivalents of oxEGCG(80%). In **c**,**d**, residues with values exceeding the mean + s.d. (red line) are colored red, and those below the mean − s.d. (blue line) are colored blue. Top, schematic representation of the αSyn regions. **e**, HPLC chromatogram of the oxEGCG(80%) reagent monitored at 220 nm. **f**, TEM image of αSyn in the presence of 10 equivalents of oxEGCG(80%). Scale bar, 300 nm.

To determine whether the observed oxEGCG interaction and directional chemical shift changes are unique to αSyn, we performed NMR titrations with ubiquitin, a stable globular protein (Supplementary Section 3). Although oxEGCG also interacted with ubiquitin (Extended Data Fig. 5a,b), the chemical shift changes occurred in multiple directions rather than mainly toward the upper-right, except for several C-terminal residues (L71, R72, R74, G75, and G76), which exhibited αSyn-like chemical shift changes. This C-terminal tail is conformationally flexible, suggesting that local structural flexibility may contribute to this behavior. Substantial chemical shift perturbations were distributed across multiple secondary-structure regions (Extended Data Fig. 5c,d), without a clear preference for a particular residue type (Extended Data Fig. 5c, bottom). The average apparent *K*_D_ was 1.9 ± 2.3 mM (Supplementary Figs. 4 and 5). Notably, oxEGCG titration induced amorphous aggregation of ubiquitin (Extended Data Fig. 5e), whereas no macroscopic precipitation was observed during the corresponding αSyn titration. Collectively, these findings demonstrate that oxEGCG can interact with both intrinsically disordered and folded proteins, but the spectral response and aggregation behavior depend on the protein.

Because laboratory-prepared oxEGCG produced chemical shift perturbations similar to those previously attributed to EGCG, we next asked whether oxidized components in EGCG preparations could account for the EGCG–αSyn interactions reported in previous NMR studies^9,22^. To address this question, we examined the interaction between αSyn and 10 equivalents of a commercially available EGCG reagent (≥80% purity; hereafter, oxEGCG(80%)), which had lower nominal purity than the high-purity EGCG (≥95% and ≥99% purity) used above. This reagent induced chemical shift perturbations in the same direction and with a residue-specific pattern similar to that induced by laboratory-prepared oxEGCG, but with markedly larger magnitudes (Fig. 2a,c,d), consistent with previous studies^9,22^. The residue-specific Δ*δ* values between free αSyn and αSyn in the presence of oxEGCG(80%) are hereafter denoted Δ*δ*^free–bound^ (Fig. 2d). Despite these large perturbations, HPLC analysis revealed a dominant EGCG peak accompanied by only minor peaks, including gallic acid (Fig. 2e). However, gallic acid induces NMR chemical shift perturbations distinct from those observed here^24^, indicating that gallic acid is unlikely to be the primary active species. Thus, the identity of the active species remains unknown. Given the time-dependent changes in EGCG oxidation products observed by HPLC (Fig. 1a), together with the smaller chemical shift changes induced by laboratory-prepared oxEGCG after 72 h of oxidation than by the commercial reagent, the active species may be transient, converting into less active forms over time. These results suggest that the EGCG–αSyn interactions reported in previous NMR studies^9,22^ may have arisen from oxidized components present in the commercial EGCG reagent. Although no macroscopic precipitation was visible in the NMR tube after addition of oxEGCG(80%), TEM revealed the formation of spherical oligomers (Fig. 2f), in agreement with a previous study^9^.

For subsequent analyses, we used oxEGCG(80%) because laboratory-prepared oxEGCG showed limited reproducibility, whereas oxEGCG(80%) yielded larger chemical shift perturbations and more reproducible NMR spectra.

### Region-dependent changes in αSyn water accessibility induced by oxEGCG(80%)

The NMR interaction analysis revealed that oxEGCG(80%) interacts with αSyn (Fig. 2a, right and 2d). To further characterize the effects of oxEGCG(80%) on αSyn using an independent NMR observable, we next performed phase-modulated clean chemical exchange experiments (CLEANEX-PM), which report on hydrogen exchange between backbone amide protons and bulk water and therefore provide information on local water accessibility upon oxEGCG(80%) binding.

Analysis of the CLEANEX-PM buildup curves revealed residue-dependent *k*_1H_ values across αSyn (Fig. 3a,b and Supplementary Figs. 6 and 7), with oxEGCG(80%) inducing both increases and decreases in *k*_1H_ at several residues (Fig. 3c). These changes indicate residue-specific increases and decreases in local water accessibility arising from oxEGCG(80%) association, changes in the αSyn conformational ensemble, or both. Furthermore, analysis of Δ*k*_1H_ by residue type showed no clear preference for particular amino acid types (Fig. 3d), suggesting that the effects of oxEGCG(80%) depend more on sequence position than on residue identity.

**Fig. 3.**
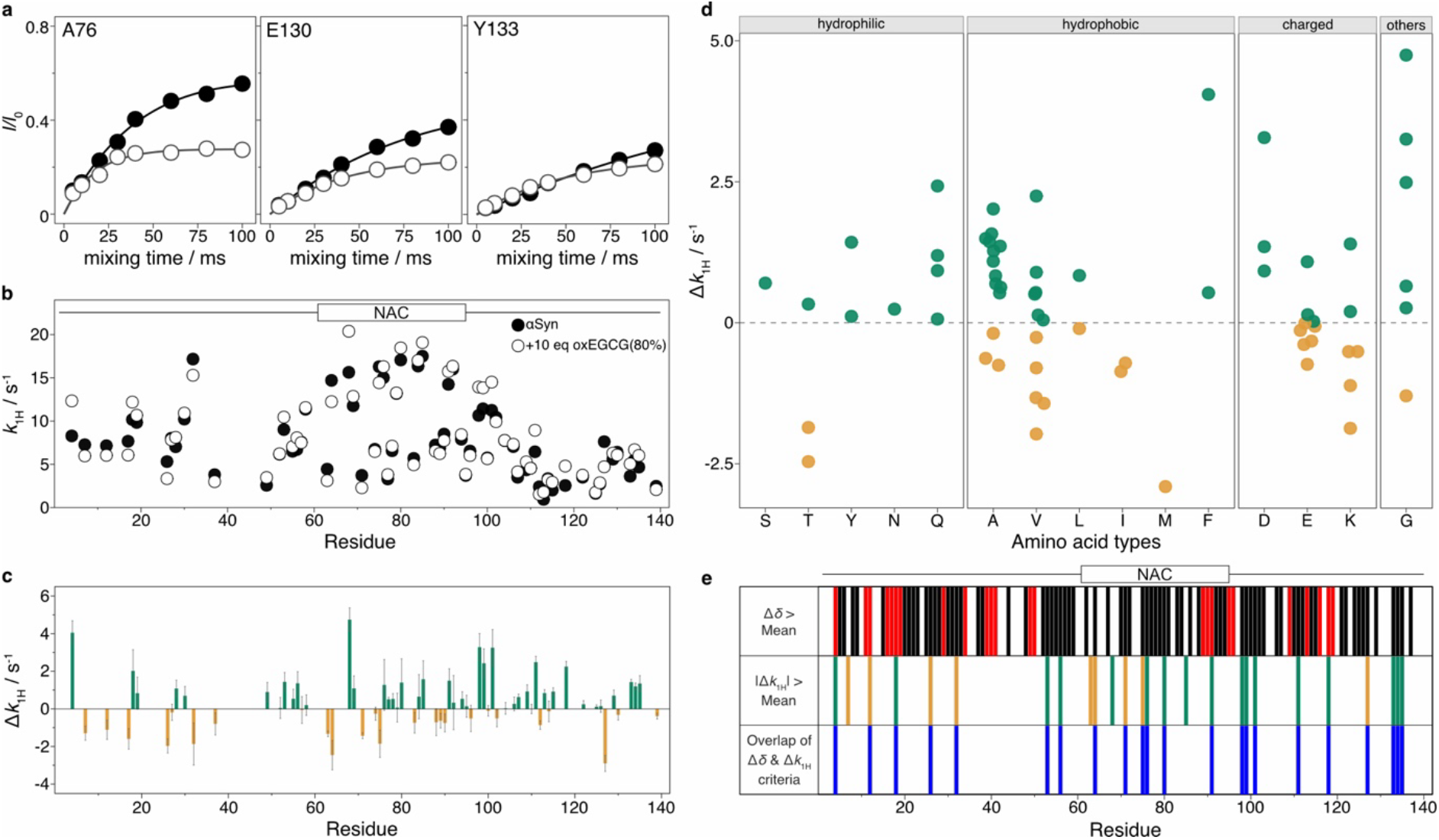
| CLEANEX-PM analysis of oxEGCG(80%)-induced changes in αSyn hydrogen exchange. **a,** Representative CLEANEX-PM buildup curves for A76, E130, and Y133 in the absence (closed circles) and presence (open circles) of oxEGCG(80%). Solid lines indicate fits to Eq. (2). **b**, Residue-specific *k*_1H_ values of αSyn in the absence (closed circles) and presence (open circles) of oxEGCG(80%). Top, schematic representation of the αSyn regions. Error bars in **a** and **b**, calculated from fit residual RMSDs and Monte Carlo–estimated s.d., respectively, are smaller than the data-point symbols. **c**, Residue-specific differences in *k*_1H_ (Δ*k*_1H_), calculated as *k*_1H_ in the presence of oxEGCG(80%) minus *k*_1H_ in its absence. Error bars indicate s.d. **d**, Scatter plot showing the distribution of Δ*k*_1H_ values for each amino acid residue type. **e**, Residue-wise comparison of Δ*δ* and |Δ*k*_1H_|. A schematic representation of the αSyn regions is shown above the plots. Top, residues with Δ*δ* values above the mean are shown in black, whereas those above the mean + s.d. are shown in red. Middle, residues with |Δ*k*_1H_| values above the mean are shown in green or orange for positive or negative Δ*k*_1H_, respectively, using the same color scheme as in **c**. Bottom, residues satisfying both Δ*δ* and |Δ*k*_1H_| criteria are indicated in blue.

We then compared the residue profiles obtained from chemical shift perturbation (Fig. 2d) and CLEANEX-PM (Fig. 3c) analyses to identify regions whose local environments were strongly perturbed in the presence of oxEGCG(80%). Residues showing both Δ*δ* and |Δ*k*_1H_| above their respective mean values were used as indicators of strong local perturbation (Fig. 3e). The selected residues were distributed throughout αSyn. Several were observed in the N-terminal region (F4, K12, A18, V26, K32, A53, A56, and T64) and the NAC region (V71, T75, A76, and K80), whereas a greater number of selected residues were clustered near the NAC– C-terminal boundary and within the adjacent C-terminal region (A91, D98, Q99, G101, G111, V118, M127, Y133, Q134, and D135). These residues are not necessarily direct binding sites, but their combined chemical shift and hydrogen exchange responses indicate that oxEGCG(80%) substantially alters their local environments. Thus, oxEGCG(80%) affects αSyn not only through broad N-terminal perturbations detected by chemical shift analysis, but also through changes in the water accessibility of selected residues near the NAC–C-terminal boundary and within the adjacent C-terminal region. The C-terminal region of αSyn is known to contribute to long-range intramolecular contacts with the N-terminal and NAC regions^25,26^. The perturbations observed near the NAC–C-terminal boundary and within the adjacent C-terminal region are therefore consistent with a change in the conformational ensemble of αSyn, potentially involving changes in these long-range intramolecular contacts.

### Shear flow modulates oxEGCG(80%)-dependent interactions with αSyn

Although the NMR interaction analysis revealed oxEGCG(80%)–αSyn interactions under static conditions (Fig. 2a, right and 2d), spontaneous αSyn fibrillation was negligible under these conditions, making it difficult to identify interactions relevant to fibrillation inhibition. We therefore used Rheo-NMR to examine oxEGCG(80%)–αSyn interactions under aggregation-promoting shear flow at an average shear rate of approximately 400 s^−1^ (Fig. 4a). After the Rheo-NMR measurement, TEM imaging showed both spherical oligomers and short, thick fibrils (Fig. 4b), despite the absence of visible precipitation in the NMR tube. The oligomers showed a broad size distribution with diameters of 15–50 nm and were larger than previously reported 10-nm αSyn oligomers^27^. The fibrils were approximately 50 nm long and 15 nm wide, making them shorter and thicker than mature αSyn fibrils, which are typically 10 nm wide and micrometers long^28^. As conventional fibril elongation proceeds primarily through monomer addition at fibril ends^17^, these morphological differences suggest that oxEGCG(80%) redirects αSyn aggregation from longitudinal fibril elongation toward surface-associated assembly, yielding larger oligomers and atypically short, thick fibrils.

**Fig. 4.**
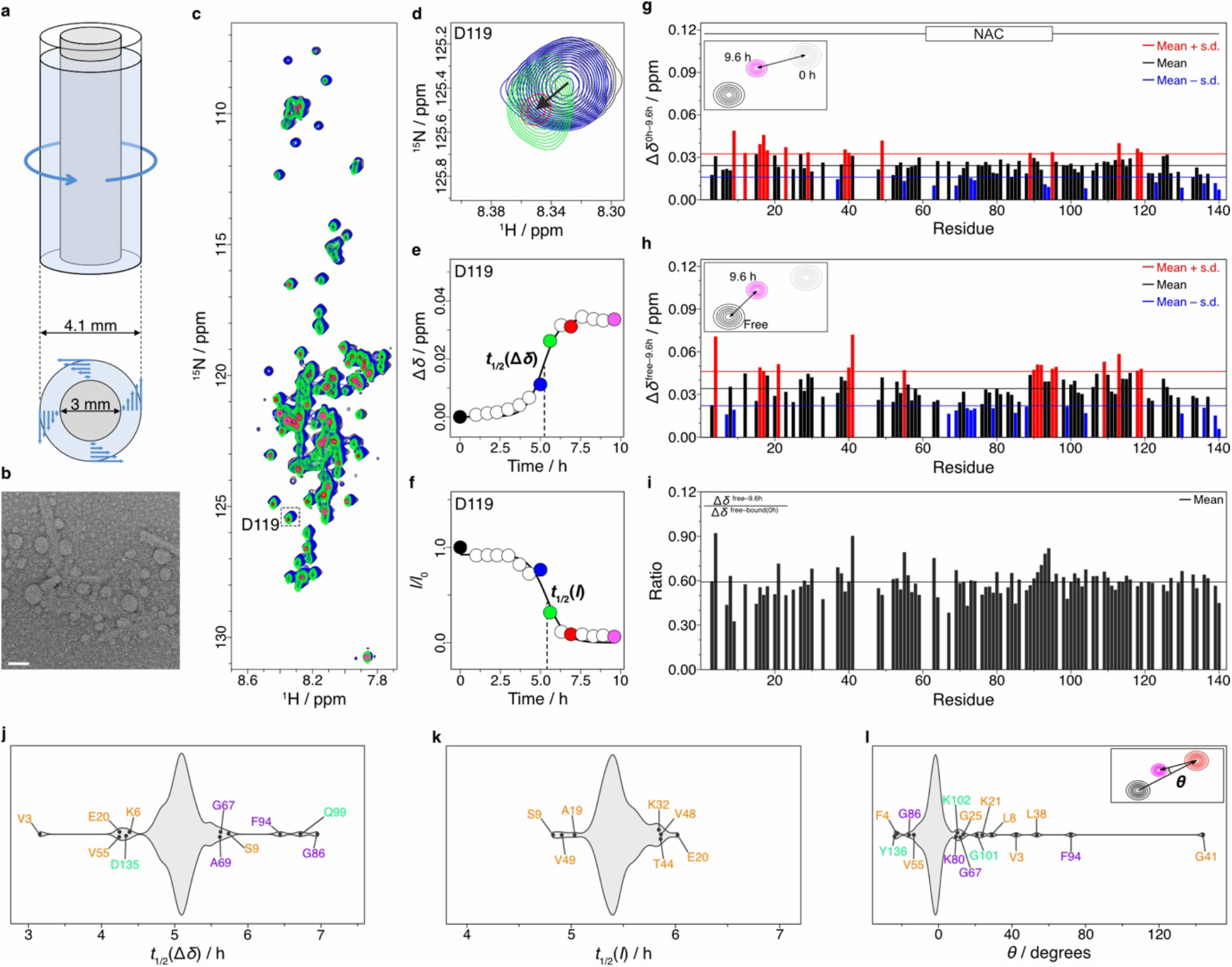
| Shear flow modulates oxEGCG(80%)-dependent perturbations in αSyn. **a**, Geometry of the sample-containing region in the Rheo-NMR setup, comprising a Shigemi tube (5 mm o.d., 4.1 mm i.d.) and a glass rod (3 mm o.d.), with a top view shown below. **b**, TEM image of αSyn aggregates after the Rheo-NMR measurement. Scale bar, 50 nm. **c,** Overlay of ^1^H-^15^N HSQC spectra of αSyn in the presence of oxEGCG(80%) at 0 (black), 5 (blue), 5.6 (green), 6.9 (red), and 9.6 (magenta) h under shear flow. **d**, Enlarged view of the spectral region for D119. **e**,**f**, Time courses of chemical shift changes (Δ*δ*) (**e**) and signal intensity decays (*I/I*_0_) (**f**) for D119. Solid lines indicate sigmoidal fits, and dashed vertical lines indicate *t*_1/2_. Colored points correspond to the spectra shown in **c**. Error bars, which represent root-mean-square deviations (RMSDs), are smaller than the data-point symbols. **g**, Residue-specific Δ*δ*^0h–9.6h^ values. The 0 h positions were recorded before shear. Residues with Δ*δ*^0h–9.6h^ values above the mean + s.d. are colored red, and those below the mean − s.d. are colored blue. Top, schematic representation of the αSyn regions. **h**, Residue-specific Δ*δ*^free–9.6h^ values between free αSyn and αSyn after 9.6 h of the Rheo-NMR measurement. Residues are colored according to mean ± s.d. thresholds as in **g**. **i**, Ratio of Δ*δ*^free–9.6h^ to Δ*δ*^free–bound^. **j**,**k**, Violin plots of *t*_1/2_ values derived from Δ*δ* (**j**) and *I/I*_0_ (**k**). **l**, Angular deviations (*θ*) between the direction from the 0 h peak positions toward those of free αSyn and the observed shear-induced movement from 0 to 9.6 h. Angles were calculated in unweighted ^1^H and ^15^N chemical shift coordinates. Only residues meeting both the apparent *K*_D_ and Rheo-NMR Δ*δ*-based *t*_1/2_ selection criteria were included. In **j–l**, outliers (below Q1 − 1.5 × IQR or above Q3 + 1.5 × IQR) are labeled and colored according to region: orange, N-terminal; purple, NAC; green, C-terminal. Inset, definition of *θ*. Q1 and Q3 denote the first and third quartiles, respectively, and IQR denotes the interquartile range.

To elucidate the molecular events underlying these morphological changes, we analyzed the time-dependent evolution of the Rheo-NMR spectra. Shear flow induced time-dependent chemical shift changes and signal intensity decays (Fig. 4c–f and Supplementary Figs. 8–12). Most signals shifted toward the lower-left, approaching the positions observed for free monomeric αSyn, suggesting partial dissociation of oxEGCG(80%) from αSyn under shear flow. Concurrently, the observed signal intensity decay was consistent with incorporation of a fraction of αSyn into oligomers or short fibrils. We quantified per-residue Δ*δ*^0h–9.6h^ values between the start (0 h) and end (9.6 h) of the Rheo-NMR measurement (Fig. 4g). Large Δ*δ*^0h–9.6h^ values were observed for many residues in the N-terminal region and for several residues in the NAC and C-terminal regions, whereas smaller Δ*δ*^0h–9.6h^ values were observed particularly near the N-terminal side of the NAC region and the C-terminal end. Residue-specific analysis revealed no strong preference for particular amino acid types (Extended Data Fig. 4b).

To assess the extent of reversion toward the free state, we calculated Δ*δ*^free–9.6h^ between free αSyn and αSyn after 9.6 h of the Rheo-NMR measurement (Fig. 4h). The analysis revealed large residual Δ*δ*^free–9.6h^ values in the N-terminal region and at several residues in the NAC and C-terminal regions, indicating that oxEGCG(80%)-dependent perturbations persisted at these sites under shear flow. To obtain a semi-quantitative estimate of the retained bound population of oxEGCG(80%) at each residue, we then calculated the ratio of Δ*δ*^free–9.6h^ (Fig. 4h) to Δ*δ*^free–bound^ (Fig. 2d). Under fast exchange conditions, the magnitude of the observed chemical shift perturbation reflects the population-weighted average of the free and bound states^29^. Thus, this ratio estimates the fraction of the initial apparent bound population that remained under shear flow. Higher values therefore indicate residues at which oxEGCG(80%) was more strongly retained under aggregation-promoting conditions. Despite the relatively large residual Δ*δ*^free–9.6h^ values in the N-terminal region (Fig. 4h), this region generally showed lower ratios, whereas residues near the NAC–C-terminal boundary (A90, A91, T92, G93, F94, and K96) showed relatively high ratios, indicating preferential retention of a subset of oxEGCG(80%)-dependent interactions at these residues under shear flow (Fig. 4i). Notably, this cluster lies near the NAC– C-terminal boundary also highlighted by the combined chemical shift and CLEANEX-PM analyses (Fig. 3e), linking preferential retention under shear to the region that showed both chemical shift perturbations and changes in water accessibility under static conditions.

To determine whether the shear-induced changes in oxEGCG(80%)–αSyn interactions were temporally coupled to αSyn incorporation into larger assemblies, we compared the kinetics of chemical shift changes and signal intensity decays. Sigmoidal fitting of chemical shift changes yielded a *t*_1/2_ of 5.1 ± 0.4 h (Fig. 4j and Supplementary Figs. 8–10), which was close to the *t*_1/2_ of 5.4 ± 0.2 h obtained for signal intensity decays (Fig. 4k and Supplementary Figs. 11 and 12). Residue-specific analysis revealed that residues identified as shorter-*t*_1/2_ outliers were predominantly located in the N-terminal region (orange), whereas longer-*t*_1/2_ outliers were mainly located in the NAC region (purple), suggesting earlier loss of oxEGCG(80%)-dependent perturbations in the N-terminal region (Fig. 4j). In contrast, the residue-specific *t*_1/2_ values for signal intensity decays showed a narrow distribution (Fig. 4k), with outliers confined to the N-terminal region, suggesting that αSyn molecules were incorporated into oligomers or short fibrils on a broadly similar timescale across most residues. The sigmoidal time courses argue against a simple first-order dissociation process driven solely by shear and instead suggest that changes in oxEGCG(80%)–αSyn interactions were coupled to assembly formation. As oligomers and short fibrillar aggregates accumulated, interactions of monomeric αSyn with these assemblies may have altered the balance of oxEGCG(80%)– αSyn interactions and contributed to the observed kinetics. Together with the TEM morphology (Fig. 4b), these results support a model in which oxEGCG(80%) favors surface-associated assembly over conventional longitudinal fibril elongation.

Next, we calculated the angular deviation (*θ*) of the shear-induced chemical shift trajectory from the reverse of the chemical shift trajectory observed upon oxEGCG(80%) addition (Fig. 4l). For most residues, *θ* values were close to zero, indicating that the dominant chemical shift change was reversion toward the free αSyn state, consistent with partial dissociation of oxEGCG(80%). However, several residues in the N-terminal region (V3, F4, L8, K21, G25, L38, G41, and V55), NAC region (G67, K80, G86, and F94), and C-terminal region (G101, K102, and Y136) exhibited larger angular deviations, indicating that their chemical shift trajectories were not explained solely by reversion to the free state. These deviations are consistent with additional shear-dependent perturbations, potentially including interactions with shear-induced assemblies.

Together, the Rheo-NMR analyses showed that shear flow partially reversed oxEGCG(80%)-dependent perturbations while promoting αSyn incorporation into larger assemblies. The perturbations that persisted near the NAC–C-terminal boundary, together with the formation of oligomers and short, thick fibrillar aggregates, indicate that oxEGCG(80%) maintains region-specific interactions and redirects αSyn aggregation under shear flow.

## Discussion

In this study, we demonstrate that the inhibition of αSyn fibrillation by EGCG is highly dependent on its oxidation state, and that the nature of oxEGCG–αSyn interactions differs between static and shear-flow conditions. ThT assays and TEM showed that EGCG oxidation is a major contributor to fibrillation inhibition and fibril remodeling, while solution NMR showed that oxidation is required for detectable αSyn binding under our NMR conditions (Figs. 1 and 2; Extended Data Fig. 2). Combined chemical shift and CLEANEX-PM analyses further identified residues with changes in both chemical shifts and water accessibility, particularly near the NAC–C-terminal boundary and within the adjacent C-terminal region (Figs. 2d and 3e). Under aggregation-promoting shear flow, many oxEGCG(80%)-dependent perturbations were partially reversed, whereas a subset of interactions near the NAC–C-terminal boundary was preferentially retained (Fig. 4). Together, these data indicate that, under shear flow, oxEGCG(80%) does not simply inhibit αSyn aggregation, but redirects it from conventional longitudinal fibril elongation toward surface-associated assembly, yielding oligomers and short, thick fibrils.

The finding that oxidized EGCG primarily accounts for these activities helps clarify the mechanism of EGCG-mediated inhibition and remodeling of amyloid assemblies (Fig. 1). Historically, the rapid autoxidation of EGCG under standard assay conditions may have been underestimated, potentially leading to the misattribution of anti-amyloidogenic properties to unoxidized EGCG. Consistent with this interpretation, previous studies have shown that oxidized EGCG can inhibit amyloid formation or remodel amyloid fibrils more strongly than unoxidized EGCG for proteins including lysozyme and amyloid-β^12,14^. By experimentally separating the unoxidized and oxidized states in our NMR analyses, we found that EGCG oxidation is a prerequisite for detectable αSyn binding under our NMR conditions and a major contributor to fibrillation inhibition and fibril remodeling (Figs. 1b–d and 2). This distinction suggests that activity in many EGCG–amyloid experiments may arise from transient oxidized EGCG species rather than the parent catechin alone.

However, establishing the oxidation dependence of EGCG activity is not sufficient to explain how oxEGCG redirects αSyn aggregation. Because amyloidogenic proteins are sensitive to mechanical agitation and shear flow, conventional solution NMR studies under static conditions cannot determine which oxEGCG–αSyn interactions persist during fibrillation. Rheo-NMR enables residue-specific monitoring of amyloid formation under controlled shear^18–21^ and, when combined with mechanistic kinetic modeling, quantitative analysis of αSyn aggregation kinetics under different solution conditions^21^. Here, we used this residue-specific capability as a dynamic filter for oxEGCG(80%)–αSyn interactions (Fig. 4). Shear flow promoted αSyn aggregation and allowed us to distinguish oxEGCG(80%)–αSyn interactions that persisted during assembly from those that weakened. Notably, the region near the NAC– C-terminal boundary highlighted by the combined chemical shift and CLEANEX-PM analyses under static conditions also showed preferential retention of oxEGCG(80%)-dependent interactions under shear (Figs. 2d, 3c, 3e and 4i). This convergence suggests that interactions in this region may contribute to the redirection of αSyn aggregation toward oligomers and short, thick fibrils under shear (Fig. 4b). A recent structural study showed that residues E83–K97 adopt distinct conformations in a lipid-associated prefibrillar αSyn oligomer and mature fibrils, highlighting the structural plasticity of this region during assembly^30^. Because the NAC region forms the structural core of αSyn amyloid fibrils^31^, preferential retention of oxEGCG(80%)-dependent interactions near the NAC–C-terminal boundary may interfere with the intermolecular packing required for conventional fibril formation and thereby favor alternative assembly pathways.

The shear-selected interaction pattern provides a basis for a redirection model analogous to molecular glue mechanisms in which small molecules enhance protein–protein interactions^32^ (Fig. 5). In this model, oxEGCG(80%) associates with αSyn under static conditions, whereas shear flow weakens many of these interactions while retaining a subset. These retained interactions may stabilize αSyn–αSyn associations, thereby favoring surface-associated assembly while limiting conventional longitudinal fibril elongation. The model provides a possible explanation for why αSyn formed spherical oligomers and short, thick fibrils in the presence of oxEGCG(80%) under shear flow, rather than typical long amyloid fibrils (Fig. 4b). Such a mechanism also helps reinterpret the apparent dual activity of EGCG as both a fibrillation inhibitor and a fibril remodeler (Fig. 1b–d). Rather than representing distinct activities, inhibition and remodeling may reflect a common oxidation-dependent ability of oxEGCG to reshape αSyn assembly pathways, redirecting aggregation toward oligomers and short fibrillar species under shear (Fig. 4b) and remodeling preformed fibrils into amorphous non-fibrillar aggregates (Fig. 1d).

**Fig. 5.**
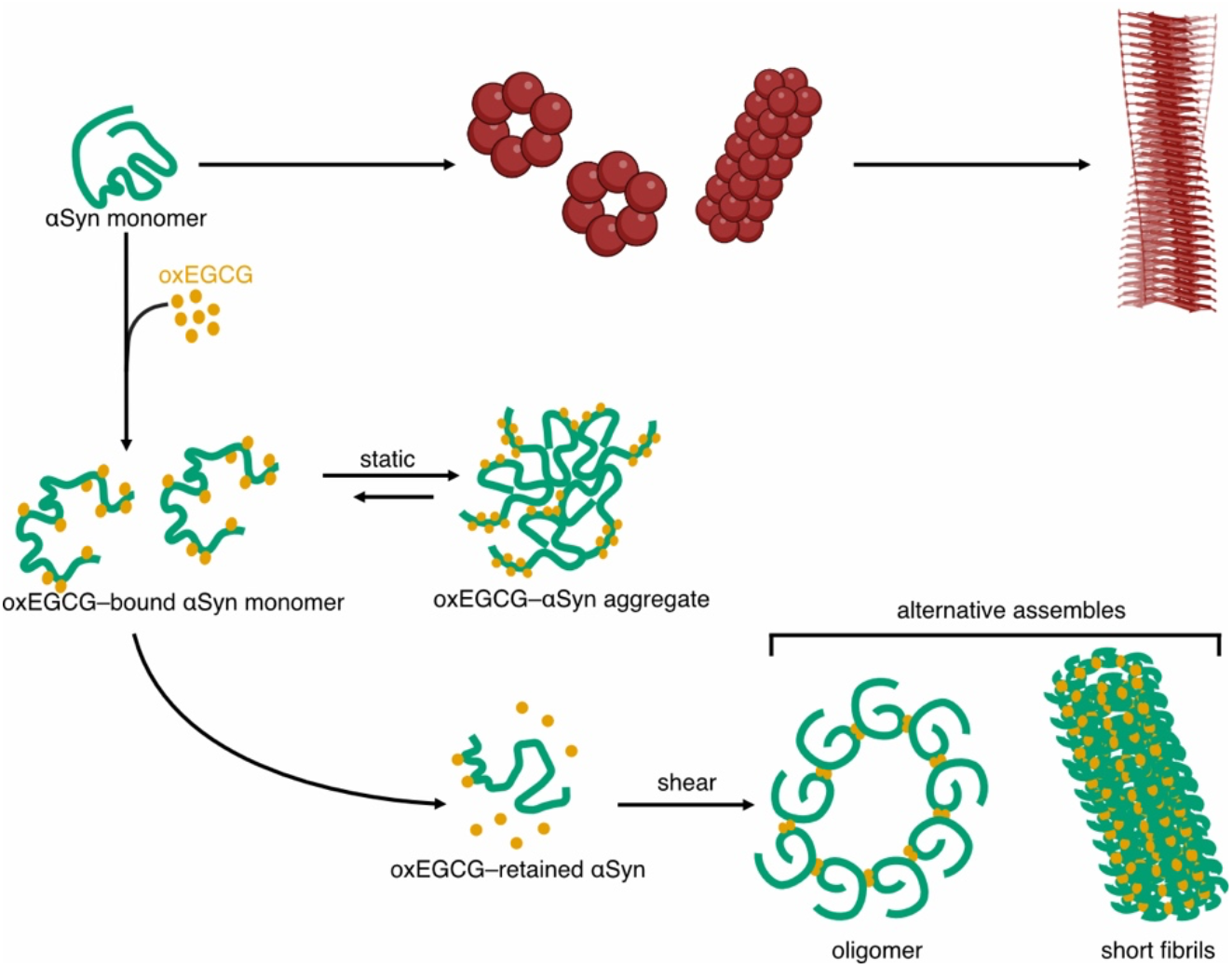
| Proposed model for oxEGCG-mediated redirection of αSyn aggregation under shear flow. OxEGCG associates with αSyn monomers and promotes oligomeric assemblies under static conditions. Under shear flow, many oxEGCG-mediated interactions are weakened, whereas retained contacts are proposed to favor alternative assemblies including oligomers and short, thick fibrils (green). In the absence of oxEGCG, shear flow promotes conventional amyloid formation, resulting in long straight fibrils (red).

Could oxidation-dependent EGCG activity also influence αSyn aggregation *in vivo*? Epidemiological studies have associated tea consumption with a reduced risk of Parkinson’s disease^7,33^, although the molecular basis for the association remains unclear. Our findings raise the possibility that locally generated oxEGCG species could influence αSyn assembly in peripheral tissues where EGCG is exposed to oxygen. One relevant site is the gastrointestinal tract, where orally ingested EGCG can accumulate in the mucosa^34^ and where the enteric nervous system has been implicated in gut-to-brain propagation models of αSyn pathology^35,36^. Although this possibility remains to be tested *in vivo*, our data provide a molecular framework for considering how EGCG oxidation could influence peripheral αSyn aggregation.

Despite these insights, the precise chemical identities of the active oxidized species remain unresolved. Determining the chemical identities and the biological properties of the shear-induced assemblies will be important next steps. Nevertheless, our findings establish oxidation state and flow conditions as key determinants of EGCG action on αSyn. OxEGCG also interacted with ubiquitin, but produced distinct chemical shift responses and aggregation behavior, indicating that the consequences of oxEGCG interactions are protein-dependent (Extended Data Fig. 5). More broadly, the effects of other amyloid inhibitors, including oxidation-prone polyphenols and small molecules that act through weak multivalent interactions, may likewise depend on their chemical state, protein context and physical environment. Accordingly, combining oxidation-controlled chemical analysis with Rheo-NMR may provide a general strategy for distinguishing direct inhibition of fibril formation from pathway redirection in amyloid-targeting small molecules.

## Materials and Methods

### Preparation of αSyn

Recombinant human [*U*-^15^N]-αSyn was expressed and purified using a previously described GST-SUMO-based protocol^37^, with modifications for isotope labeling. Briefly, GST-SUMO-tagged αSyn was expressed in *E. coli* BL21(DE3) cells using a pET23a vector. The cells were grown overnight at 37 °C in 200 mL M9 medium containing 50 µg/mL ampicillin and 0.5 g/L ^15^NH_4_Cl (Cambridge Isotope Laboratories) as the sole nitrogen source. The resulting culture was used to inoculate 2 L M9 medium. At an optical density at 600 nm (OD_600_) of approximately 0.6, protein expression was induced with 1 mM IPTG, followed by incubation at 25 °C for 15 h. The cells were harvested by centrifugation, resuspended in 50 mM Tris-HCl buffer (pH 7.5) containing 1 mM EDTA, 1 mM DTT, and one cOmplete protease inhibitor cocktail tablet (Roche), and lysed by sonication. The lysate was clarified by centrifugation, and the filtered supernatant was applied to a Glutathione Sepharose 4B column (Cytiva). After washing, GST-SUMO-tagged αSyn was eluted with reduced glutathione and cleaved with GST-SENP2 overnight at 4 °C during dialysis. The cleaved sample was purified by anion-exchange chromatography on a HiTrap Q HP column (Cytiva) and then passed through a Glutathione Sepharose 4B column to remove the cleaved GST-SUMO tag and GST-SENP2. The flow-through containing tag-free αSyn was dialyzed against ultrapure water, lyophilized, and stored at 4 °C until use.

### HPLC analysis

EGCG (≥99% purity; Taiyo Kagaku) was dissolved in ultrapure water at 5 mg/mL and stirred at 25 °C, with samples collected at 0, 2, 4, 6, 8, 16, 24, 32, 40, and 48 h. The samples were diluted with ultrapure water to 500 ng/µL, and 5-µL aliquots were analyzed using a Prominence HPLC system equipped with an SPD-M40 PDA detector (SHIMADZU), and chromatographic separation was performed on a CAPCELL C18 column (SG300Å; 5 µm; 4.6 mm × 250 mm; SHISEIDO) at 30 °C. The flow rate was 0.8 mL/min, with detection at 220 nm. The mobile phase consisted of (A) ultrapure water containing 0.5% (v/v) trifluoroacetic acid (TFA) and (B) acetonitrile containing 0.5% (v/v) TFA. The gradient program was as follows: 12% B for 3 min, 12–25% B over 28 min, 25–95% B over 1 min, 95% B for 3 min, 95–12% B over 3 min, followed by equilibration at 12% B for 10 min.

### Thioflavin T fluorescence assay

The effects of EGCG and oxEGCG on αSyn amyloid fibrillation were evaluated using a thioflavin T (ThT) fluorescence assay. OxEGCG was prepared by stirring EGCG at 6 mg/mL in 50 mM Tris-HCl buffer (pH 7.4) containing 150 mM NaCl at 25 °C for 72 h. Fibrillation assays were performed in a sealed 96-well plate (Greiner Bio-One) at 50 µL per well. Each mixture contained 500 µM αSyn and 10 µM ThT in the same buffer. The following five conditions were tested: no additive, 1 mM EGCG, 1 mM EGCG with 1 mM DTT, oxEGCG corresponding to an initial EGCG concentration of 1 mM, and 1 mM DTT. Because oxEGCG comprised a heterogeneous mixture of oxidation products, its concentration was expressed as the initial EGCG concentration. The plate was incubated at 37 °C in an SH-9500N microplate reader (CORONA ELECTRIC) with cyclic agitation at 1100 rpm, and ThT fluorescence intensity was measured at 15-min intervals with excitation at 450 nm and emission at 485 nm.

### Amyloid fibril remodeling analysis

The effects of EGCG and oxEGCG on preformed αSyn amyloid fibrils were examined using the same buffer, additive concentrations, temperature and agitation conditions as in the ThT fluorescence assay, but without ThT and with preformed fibrils instead of monomeric αSyn. Preformed αSyn fibrils were prepared by incubating 500 µM αSyn in 50 mM Tris-HCl buffer (pH 7.4) containing 150 mM NaCl with continuous shaking for 72 h at 25 °C. After incubation with cyclic agitation at 1100 rpm for 24 h at 37 °C, fibril remodeling was assessed by TEM.

### TEM imaging

Protein aggregates were adsorbed onto plasma-cleaned 400-mesh copper grids bearing a carbon-coated collodion film, negatively stained with 1% EM Stainer (Nisshin EM), and imaged using an H-7650 transmission electron microscope (Hitachi High-Technologies).

### NMR interaction experiments

[*U*-^15^N]-αSyn and [*U*-^15^N]-ubiquitin were each prepared at 200 µM in 40 mM MOPS buffer (pH 7.0) containing 150 mM NaCl and 5% D_2_O. The oxEGCG sample was prepared as described for the ThT fluorescence assay. Three sets of NMR interaction experiments were performed at 25 °C on an Avance II 700-MHz spectrometer equipped with a 5-mm TCI triple-resonance cryogenic probe (Bruker). For all experiments, 2D ^1^H-^15^N HSQC spectra were recorded with 512 × 128 complex points. First, the interaction between αSyn and 8 molar equivalents of EGCG (≥95% purity) was examined in the presence of 10 molar equivalents of DTT. Second, oxEGCG was titrated into αSyn at 0.5, 1, 2, 4, 8, and 16 molar equivalents and into ubiquitin at 0.5, 1, 2, 4, and 8 molar equivalents. Third, the interaction between αSyn and 10 molar equivalents of oxEGCG(80%) (Sigma-Aldrich) was examined. NMR data were processed using TopSpin (Bruker), and signal intensities and chemical shifts were analyzed using PINT^38^. Resonance assignments for αSyn and ubiquitin were taken from our previous study^39^. Weighted chemical shift differences (Δ*δ*) were calculated as:

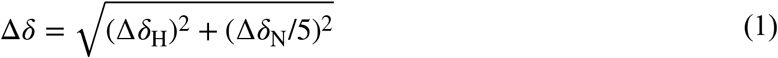

where Δ*δ*_H_ and Δ*δ*_N_ represent the chemical shift differences in the ^1^H and ^15^N dimensions, respectively.

### CLEANEX-PM analysis

[*U*-^15^N]-αSyn samples were prepared at 500 μM in 40 mM MOPS buffer (pH 7.0) containing 150 mM NaCl and 5% D_2_O, either alone or in the presence of 5 mM oxEGCG(80%). 2D CLEANEX-PM spectra with 1024 × 128 complex points were recorded at 25 °C on an 800-MHz Avance III spectrometer equipped with a 5-mm TCI triple-resonance cryogenic probe (Bruker). Mixing times (*τ*_m_) were set to 5, 10, 20, 30, 40, 60, 80, and 100 ms. The water-amide proton exchange rates (*k*_1H_) were estimated by fitting the CLEANEX-PM data to the following equation using the nlsLM function in the minpack.lm package^40^:

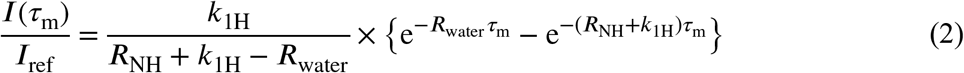

where *I*(*τ*_m_)*/I*_ref_ represents the normalized CLEANEX-PM intensity, *I*(*τ*_m_) is the peak intensity at each mixing time, *I*_ref_ is the peak intensity in the reference spectrum, and *R*_NH_ and *R*_water_ denote the effective relaxation rates of the amide proton and bulk water during the CLEANEX-PM mixing period, respectively^41^. During fitting, all parameters were initially fitted for each residue, and *R*_water_ was then set to the median of its fitted values for global fitting across residues to optimize *R*_NH_ and *k*_1H_. For comparison, the same data were fitted to a zero-intercept linear model with the lm function in R (version 4.4.2). The two models were compared using an *F*-test, with *p* < 0.05 used to select the CLEANEX-PM model over the linear model. Residues with poor fits, defined as residual sum of squares (RSS) values greater than the mean RSS plus one s.d. across all fitted residues, were excluded from further analysis. The standard deviations of the fitted *k*_1H_ values were estimated by Monte Carlo error analysis with 100 iterations. Residue-wise comparisons between conditions used residues retained in both datasets after filtering. For the analysis of Δ*k*_1H_ by amino acid type, residues were retained only when the relative standard error of *k*_1H_ obtained from nonlinear fitting was ≤ 0.5 in both conditions. This cutoff was used to exclude poorly determined exchange rates by requiring each estimate to be at least twice its standard error.

### Rheo-NMR measurements

A [*U*-^15^N]-αSyn sample was prepared at 500 µM in 40 mM MOPS buffer (pH 7.0) containing 150 mM NaCl and 5% D_2_O in the presence of 5 mM oxEGCG(80%). Rheo-NMR experiments were performed at 25 °C using a µRheo-NMR apparatus (AirPro)^42,43^ installed on an Avance I 600-MHz spectrometer equipped with a 5-mm TCI triple-resonance cryogenic probe (Bruker). 2D ^1^H-^15^N HSQC spectra were continuously acquired with 512 × 128 complex points and 8 scans under both static and sheared conditions. The spinning rate was set to 20 Hz, corresponding to an average shear rate of approximately 400 s^−1^, to accelerate αSyn aggregation and enable analysis within approximately 10 h. Signal intensities were normalized to their initial values (*I*/*I*_0_), and Δ*δ* values were calculated as described above.

### Analysis of chemical shift differences and intensity changes

Apparent dissociation constants (*K*_D_) were obtained by nonlinear least-squares fitting of the titration Δ*δ* values using the nlsLM function in R (version 4.4.2). The data were fitted to a 1:1 binding model assuming fast exchange on the chemical shift timescale:

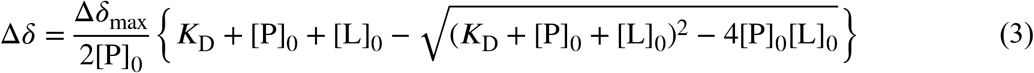

where [P]_0_ is the total protein concentration, [L]_0_ is the total EGCG-equivalent concentration of oxEGCG, and Δ*δ*_max_ is the maximum chemical shift change at saturation^29^. For each dataset, fits with RSS exceeding the mean RSS plus one s.d. across successfully fitted residues were excluded. To reduce the influence of variation in residue-specific *K*_D_ estimates on the arithmetic mean *K*_D_ value^44^, an additional empirical filter was applied by excluding estimates outside the mean ± one s.d., calculated from the RSS-retained estimates. The remaining apparent *K*_D_ values were summarized as the mean ± s.d. across residues.

Time-dependent Δ*δ* values and normalized signal intensities obtained from Rheo-NMR measurements were fitted to the following equation by nonlinear least squares using the same procedure:

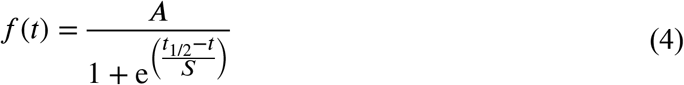

where *f*(*t*) represents the fitted Δ*δ* value or normalized signal intensity at time *t* under shear, *A* is the upper asymptotic value of the fitted curve, *t*_1/2_ is the midpoint of the fitted transition at which *f*(t) = *A*/2, and *S* is the scale parameter that determines the steepness of the transition. For each dataset, residues with poor fits, defined as RSS values greater than the mean RSS plus one s.d. across all fitted residues, were excluded from further analysis. Following RSS-based fit screening, an empirical upper cutoff of the mean *t*_1/2_ plus one standard deviation was applied to *t*_1/2_ values obtained from chemical shift changes across residues. This cutoff excluded only residues whose fitted transition midpoints lay beyond the 9.6 h observation window.

## Data availability

All data supporting the findings of this study are available within the article, its Supplementary Information and the accompanying Source Data. Additional information is available from the corresponding author upon reasonable request.

## Code availability

Custom R scripts used to analyze ThT fluorescence, NMR titration, Rheo-NMR, and CLEANEX-PM data are provided as Supplementary Software.

## Acknowledgements

This work was supported by JSPS KAKENHI Grants (21H02442, 22H04842, 22H05088, 25K00268, and 25K02240), the Scholar and Joint Research Scholar Programs of the Toyota Physical and Chemical Research Institute, research grants from the Japan Keirin Autorace Foundation (2021M-224 and 2023M-363), the Toyo Suisan Foundation, the Research Institute for Production Development, the Mitsubishi Foundation, the Hachiro Honjo Ocha Foundation, the Takeda Science Foundation, and the Collaborative Research Program of Institute for Protein Research, Osaka University (CR-26-02 and NMRCR-26-02). TEM observations were performed at the North Campus Instrumental Analysis Station, Kyoto University. We thank Taiyo Kagaku for providing high-purity EGCG (≥99% purity).

## Author contributions

C.M.Y., R.Y., K.U., Y. Shimada, Y. Shuto, and S.S. performed experiments and analyzed data. D.M., H.O., M.S., and A.F. provided experimental guidance and contributed to data interpretation and discussion. C.M.Y. wrote the initial draft of the manuscript. M.S. contributed to manuscript writing and revision. K.S. conceived and designed the study, acquired funding, supervised the project, contributed to data interpretation and discussion, and wrote and finalized the manuscript. All authors reviewed and approved the final manuscript.

