## Extended Data for "Oxidized EGCG redirects α-synuclein aggregation through shear-selected interactions"

### **Table of Contents**

Extended Data Figs. 1–5

Extended Data Table 1

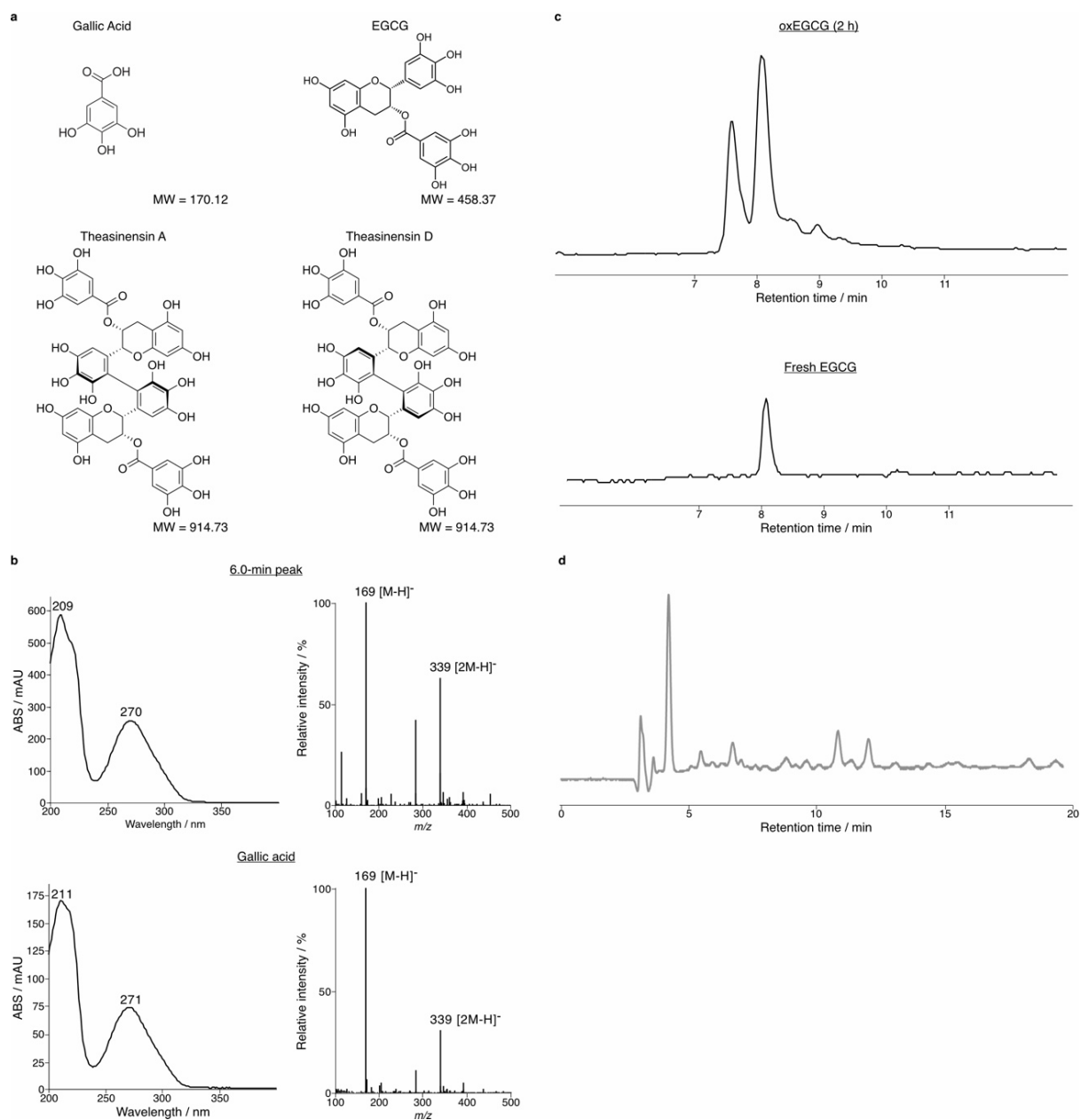

**Extended Data Fig. 1 | LC-MS and HPLC analyses of oxidized EGCG products.** **a**, Chemical structures of EGCG, gallic acid, theasinensin A, and theasinensin D. **b**, UV-Vis and mass spectra of the 6-min HPLC fraction and a gallic acid standard. **c**, Negative-ion SIM chromatograms at $m/z$ 913 ($[M-H]^-$) of oxEGCG and freshly prepared EGCG. **d**, HPLC chromatogram of the sample containing EGCG and DTT after the ThT fluorescence assay.

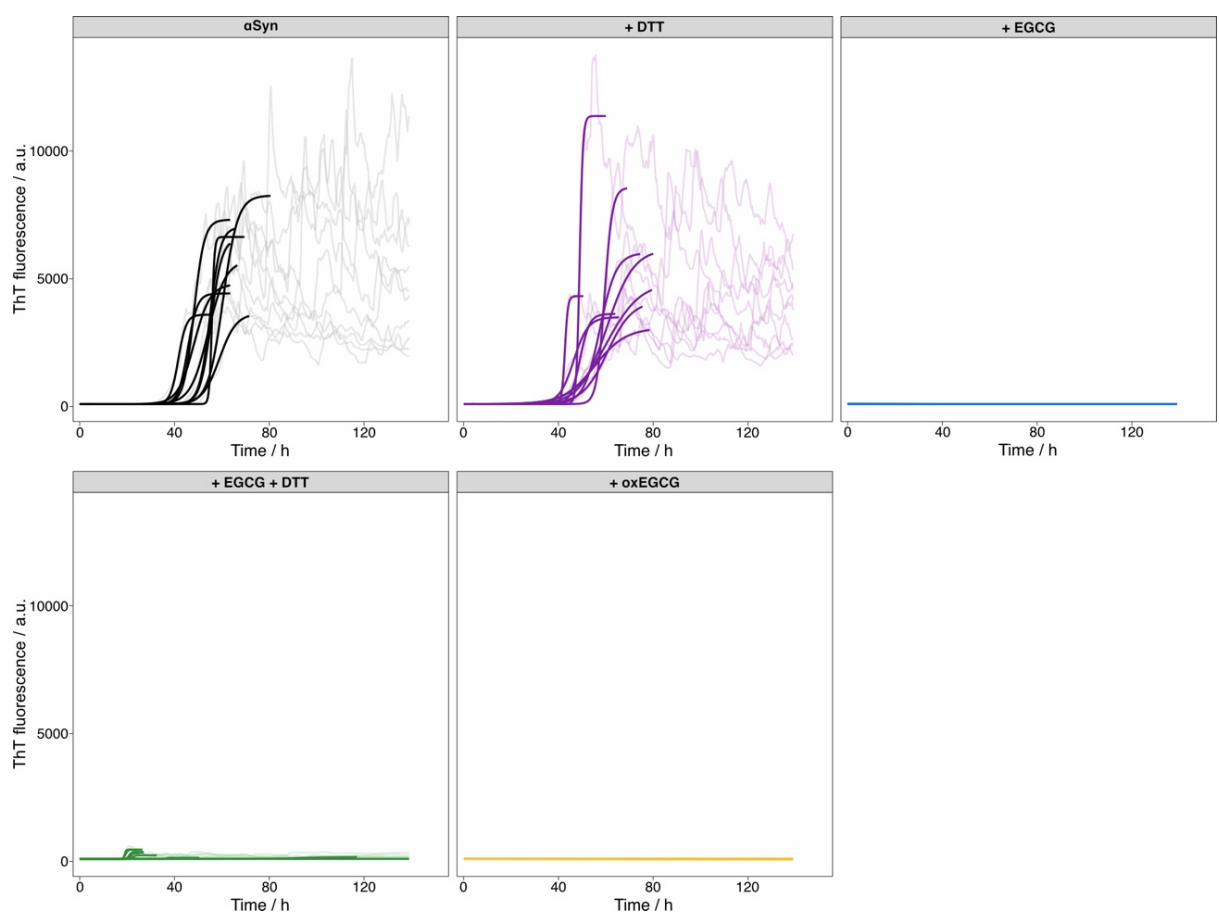

**Extended Data Fig. 2 | Individual fitting of ThT traces.** Individual ThT fluorescence traces for each condition with the corresponding fitted curves within the selected fitting interval. Raw fluorescence traces are shown in light colors and fitted curves in solid colors.

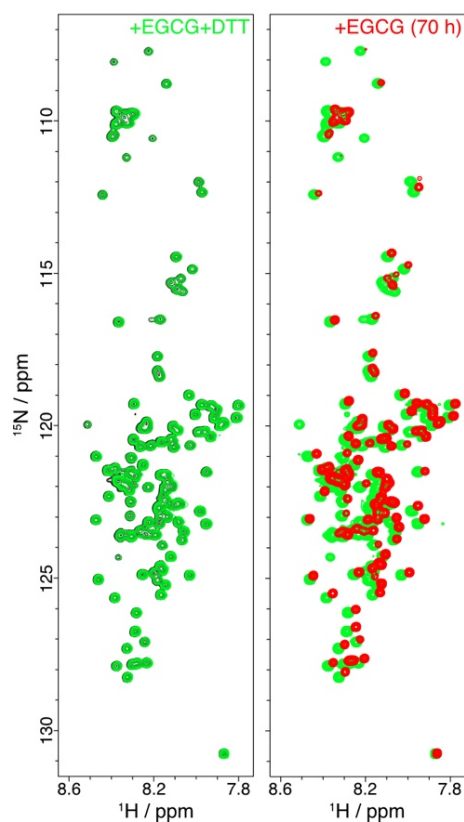

**Extended Data Fig. 3** | NMR analysis of the interaction between αSyn and unoxidized EGCG. Left, overlay of $^1\text{H}$-$^{15}\text{N}$ HSQC spectra of αSyn alone (black) and in the presence of 8 equivalents of high-purity EGCG ($\geq 95\%$ purity) supplemented with DTT (green). No detectable chemical shift changes were observed under these conditions. Right, overlay of the same EGCG-containing spectrum shown in the left panel (green) and the spectrum recorded after 70 h (red). Detectable chemical shift changes appeared after 70 h, consistent with gradual autoxidation of EGCG.

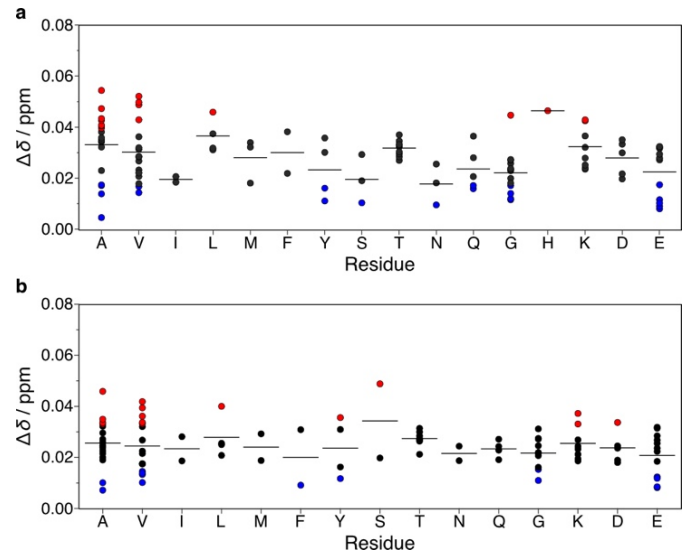

**Extended Data Fig. 4 | Residue-type analysis of αSyn chemical shift changes.** **a**, Weighted chemical shift differences ($\Delta\delta$) in the presence of 8 equivalents of laboratory-prepared oxEGCG, grouped by amino acid type. **b**, $\Delta\delta$ between 0 and 9.6 h of the Rheo-NMR measurement in the presence of oxEGCG(80%), grouped by amino acid type. The same color schemes as in Figs. 2c and 4g are used in **a** and **b**, respectively. Short horizontal lines indicate the mean $\Delta\delta$ for each amino acid type.

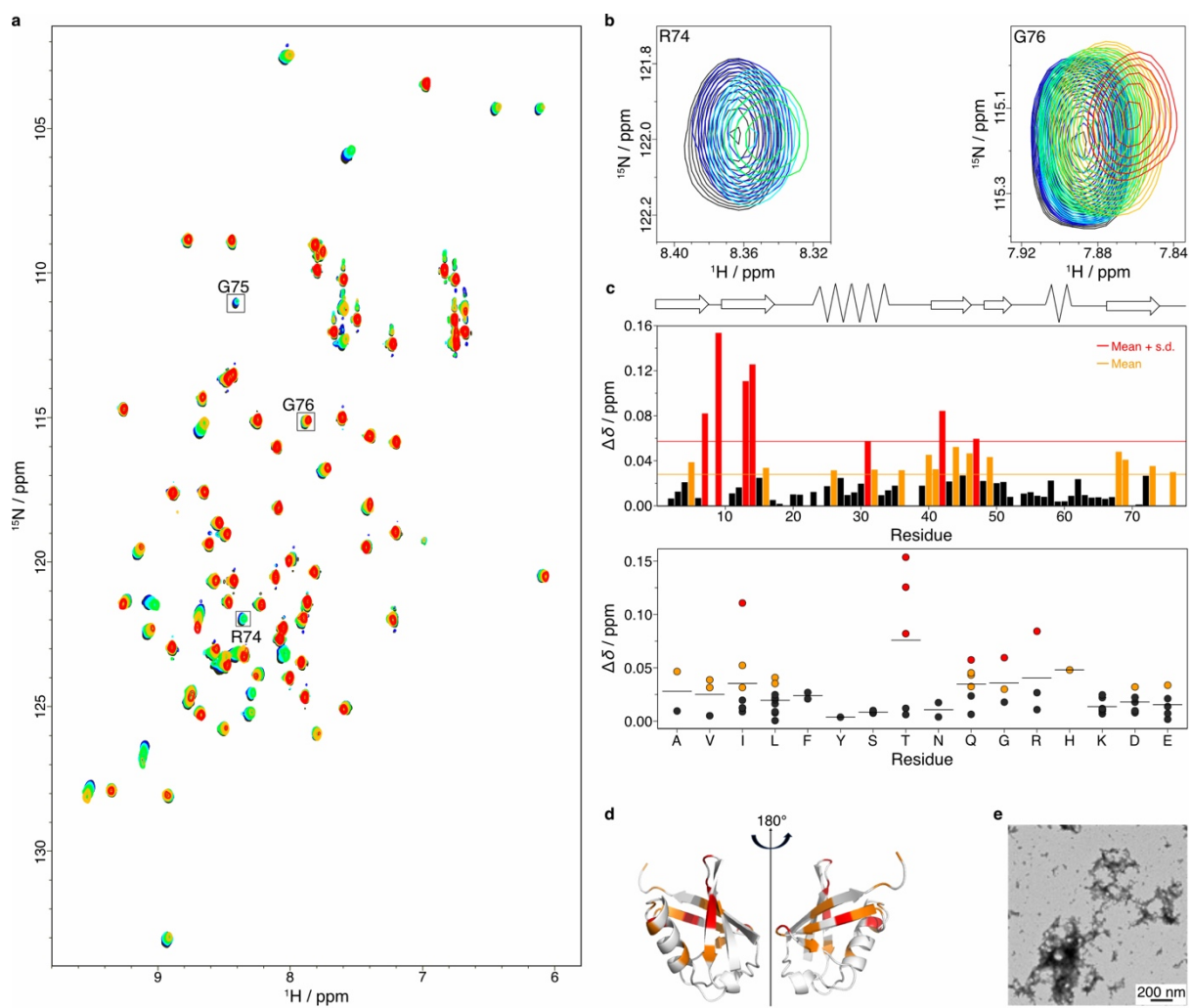

**Extended Data Fig. 5 | NMR titration of ubiquitin with oxEGCG.** **a**, Overlay of $^1\text{H}$-$^{15}\text{N}$ HSQC spectra of ubiquitin in the presence of laboratory-prepared oxEGCG at 0 (black), 0.5 (blue), 1 (cyan), 2 (green), 4 (yellow), and 8 (red) equivalents. Residues R74, G75, and G76 are labeled to highlight signals that shift toward the upper right upon addition of laboratory-prepared oxEGCG. **b**, Enlarged views of representative spectral regions for R74 and G76. **c**, Residue-specific weighted chemical shift differences ($\Delta\delta$) of ubiquitin in the presence of 8 equivalents of laboratory-prepared oxEGCG. Residues with $\Delta\delta$ exceeding the mean are colored orange, and those exceeding the mean + s.d. are colored red. Top, schematic representation of ubiquitin secondary structure (wavy lines, α-helices; arrows, β-strands). Bottom, residue-type analysis of ubiquitin $\Delta\delta$ values grouped by amino acid type, with horizontal lines indicating the mean $\Delta\delta$ for each residue type. **d**, $\Delta\delta$ values mapped onto the ubiquitin structure (PDB accession code: 1UBQ) using the color scheme in (c). **e**, TEM image of ubiquitin aggregates formed after titration with laboratory-prepared oxEGCG. Scale bar, 200 nm.

**Extended Data Table 1** | Model selection and fitted parameters for ThT fluorescence traces.

| Condition | Selected model | $n$ | $t_{1/2}$ (h) | $S$ (h) | A (a.u.) | Slope (a.u./h) |
| --- | --- | --- | --- | --- | --- | --- |
| αSyn | Logistic | 10 | $52.4 \pm 6.1$ | $2.5 \pm 1.0$ | $5796 \pm 1616$ | – |
| +DTT | Logistic | 10 | $54.1 \pm 6.7$ | $3.6 \pm 2.4$ | $5561 \pm 2614$ | – |
| +EGCG | Linear | 10 | – | – | – | $0.0 \pm 0.0$ |
| +EGCG+DTT | Logistic | 5 | $20.4 \pm 1.1$ | $0.4 \pm 0.1$ | $395 \pm 65$ | – |
| | Linear | 1 | – | – | – | $0.0 \pm 0.0$ |
| +oxEGCG | Linear | 9 | – | – | – | $0.0 \pm 0.0$ |

Values are mean ± s.d.; $n$ indicates the number of traces assigned to each model.
