## Supplementary Info for "Oxidized EGCG redirects α-synuclein aggregation through shear-selected interactions"

### **Table of Contents**

1. LC-MS and HPLC analyses of oxidized EGCG products

2. Analysis of thioflavin T fluorescence intensities

3. Preparation of ubiquitin

Supplementary Figs. 1–12

Supplementary References

### 1. LC-MS and HPLC analyses of oxidized EGCG products

EGCG has been reported to form various oxidation products, including gallic acid and theasinensins A and D (Extended Data Fig. 1a)<sup>1,2</sup>. In our HPLC analysis, the EGCG peak at ~13 min decreased over time, whereas a peak at ~6 min increased (Fig. 1a). The 1-mL fraction eluting at ~6 min was collected and concentrated using a CVE-2000 centrifugal evaporator (Tokyo Rikakikai). The dried sample was redissolved in 50 μL of 50% (v/v) acetonitrile, and a 3-μL aliquot was analyzed by LC-PDA-MS. Gallic acid was identified by comparison with a standard (Extended Data Fig. 1b).

For LC-MS detection of EGCG oxidation products, EGCG was dissolved at 5 mg/mL in 200 mM Tris-HCl buffer (pH 8.0) and stirred at 37 °C for 2 h. The resulting oxEGCG solution was diluted to 500 ng/μL with 50% (v/v) acetonitrile and centrifuged. The supernatant was applied to a MonoSpin C18 column (GL Sciences), and the flow-through was further diluted to 50 ng/μL with 50% (v/v) acetonitrile. A 99-μL aliquot of the diluted sample was mixed with 1 μL of 1 μg/μL 3,3',4',7-tetrahydroxyflavone (FUJIFILM Wako Pure Chemical) as an internal standard. A 1-μL aliquot of the mixture was analyzed using a Prominence HPLC system equipped with an SPD-M20A diode array detector and coupled to an LCMS-2020 mass spectrometer with electrospray ionization (SHIMADZU). Chromatographic separation was performed on a Mightysil RP-18 GP column (Kanto Chemical) at 40 °C with a flow rate of 0.2 mL/min. The mobile phase consisted of (A) ultrapure water containing 0.1% (v/v) formic acid and (B) acetonitrile containing 0.1% (v/v) formic acid. The gradient program was 1% B for 2 min, followed by a linear increase to 50% B over 10 min. To detect theasinensins A and D, negative-ion selected ion monitoring (SIM) was performed at $m/z$ 913, corresponding to the calculated $[M-H]^-$ ion ($C_{44}H_{33}O_{22}^-$, 913.1469). The MS parameters were as follows: nebulizer gas flow, 1.5 L/min; drying gas flow, 15 L/min; detector voltage, 1.8 kV; heat block temperature, 200 °C; and desolvation line temperature, 250 °C.

The negative-ion SIM chromatogram of oxEGCG showed two peaks at $m/z$ 913 (Extended Data Fig. 1c), consistent with the presence of theasinensins A and D. Their identities could not be confirmed because of their low abundance, although both compounds have previously been reported as EGCG oxidation products<sup>2</sup>. Freshly prepared EGCG also showed a small peak at $m/z$ 913, but only a single peak was detected (Extended Data Fig. 1c), suggesting that trace amounts of these species were initially present or formed during sample preparation or analysis.

HPLC analysis was also performed immediately after the ThT fluorescence assay (Fig. 1b) on the sample containing EGCG and DTT. The EGCG peak at ~13 min was barely detectable, indicating extensive EGCG oxidation during the assay (Extended Data Fig. 1d).

### 2. Analysis of Thioflavin T fluorescence intensities

Each ThT fluorescence trace recorded at 15-min intervals was smoothed by replacing each data point with the mean within a centered 2-h window (8 data points). The smoothed traces were fitted independently to both a logistic model and a linear model using R (version 4.4.2) in RStudio. Traces for which neither model converged were excluded. The logistic model was defined as:

$$I(t) = I_{\text{base}} + \frac{A - I_{\text{base}}}{1 + e^{\left(\frac{t_{1/2} - t}{S}\right)}}$$

where $I(t)$ represents the fitted ThT fluorescence intensity at time $t$, $I_{\text{base}}$ is the lower asymptotic value fixed at 100 a.u., approximately corresponding to the mean fluorescence across all conditions during the first 10 h, $A$ is the upper asymptotic value of the fitted curve, $t_{1/2}$ is the midpoint of the fitted transition at which $I(t) = (A + I_{\text{base}})/2$, and $S$ is the scale parameter that determines the steepness of the transition. For the logistic fits, fluorescence values below 100 a.u. were set to 100 a.u. before fitting. To determine whether each trace was better described by the logistic or linear model, the corrected Akaike Information Criterion (AICc) was compared between the two models. The logistic model was selected when $\Delta\text{AICc} = \text{AICc}_{\text{logistic}} - \text{AICc}_{\text{linear}} < -2$; otherwise, the linear model was selected (Extended Data Fig. 2; Extended Data Table 1). Traces for which the logistic model was selected were considered to show detectable ThT-positive fibrillation if the fitted plateau exceeded 300 a.u.; this threshold was defined as the mean fluorescence across all conditions during the first 10 h plus three s.d. Traces with fitted plateaus below this threshold were excluded from the calculation of the fibrillation kinetic parameters. For each condition, the parameters of the selected model were averaged across traces, and the mean parameters were used to generate a representative fitted curve.

#### 3. Preparation of ubiquitin

Recombinant human [$U$-$^{15}\text{N}$]-ubiquitin was expressed in *E. coli* BL21(DE3) cells using a pET47b vector. The cells were grown overnight at 37 °C in 200 mL M9 medium containing 20 µg/mL kanamycin and 0.5 g/L $^{15}\text{NH}_4\text{Cl}$ as the sole nitrogen source. The resulting culture was used to inoculate 2 L M9 medium. At an optical density at 600 nm ($\text{OD}_{600}$) of approximately 0.6, protein expression was induced with 1 mM IPTG, followed by incubation at 25 °C for 15 h. The cells were harvested by centrifugation, resuspended in 10 mL of 50 mM sodium acetate buffer (pH 4.5), and lysed by sonication. The lysate was heated at 85 °C for 20 min and cooled to 25 °C. After centrifugation, the supernatant was filtered through a 0.45-µm membrane and purified by cation-exchange chromatography on a HiTrap SP HP column (Cytiva) using a 150–250 mM NaCl gradient in 50 mM sodium acetate buffer (pH 4.5). The eluate was further purified by size-exclusion chromatography on a Superdex 75 pg 16/60 column (Cytiva) in 25 mM Tris-HCl buffer (pH 8.0) containing 150 mM NaCl. Fractions containing ubiquitin were pooled, lyophilized, and stored at 4 °C until use.

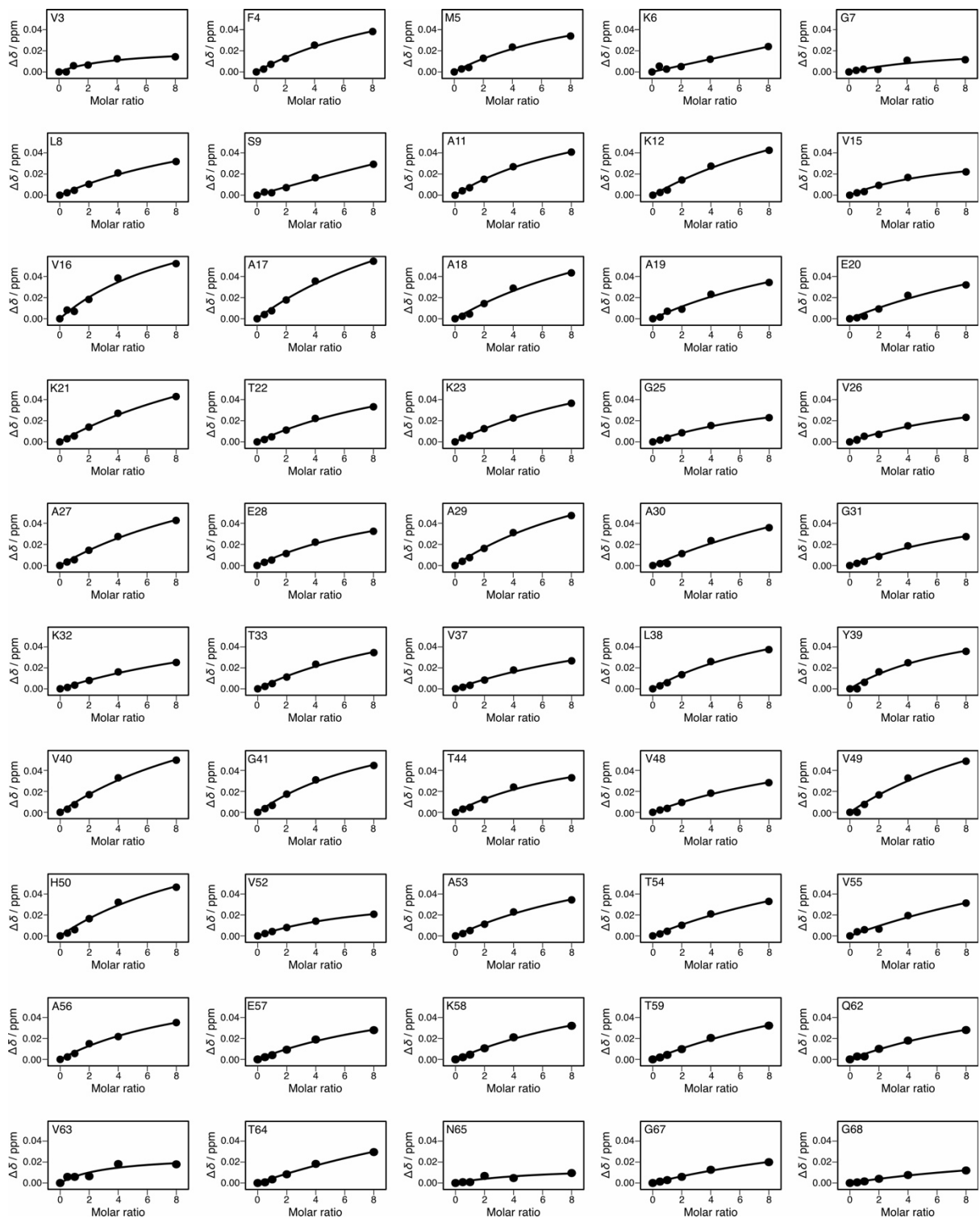

**Supplementary Fig. 1 | Titration curves of chemical shift changes for αSyn residues V3–G68 upon addition of oxEGCG.** Points represent residue-specific weighted chemical shift differences ($\Delta\delta$), and solid lines indicate fits to a 1:1 binding model used to estimate apparent $K_D$ values. Error bars, which represent root-mean-square deviations (RMSDs), are smaller than the data-point symbols.

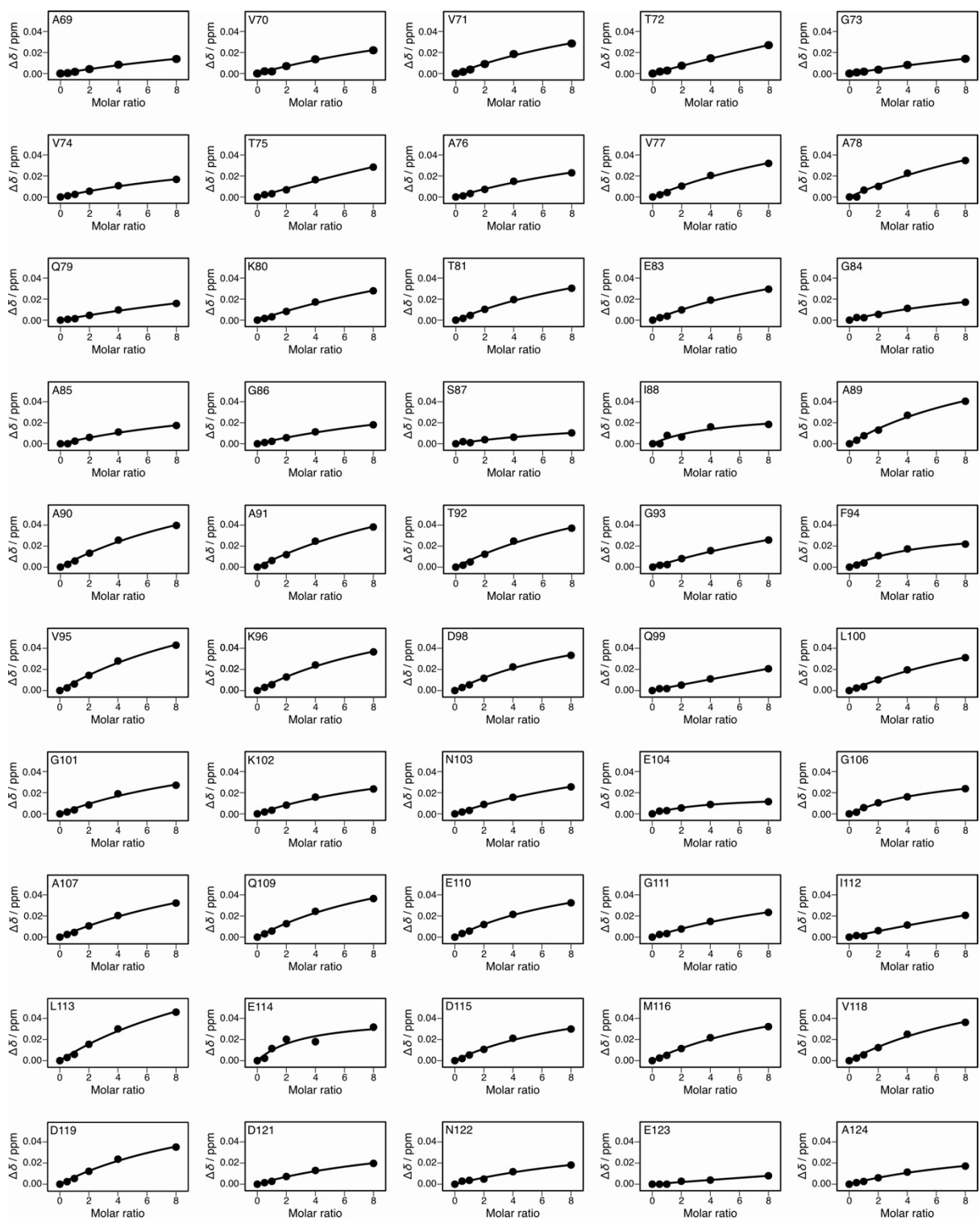

**Supplementary Fig. 2 | Titration curves of chemical shift changes for αSyn residues A69–A124 upon addition of oxEGCG.** Points represent residue-specific weighted chemical shift differences ($\Delta\delta$), and solid lines indicate fits to a 1:1 binding model used to estimate apparent $K_D$ values. Error bars, which represent root-mean-square deviations (RMSDs), are smaller than the data-point symbols.

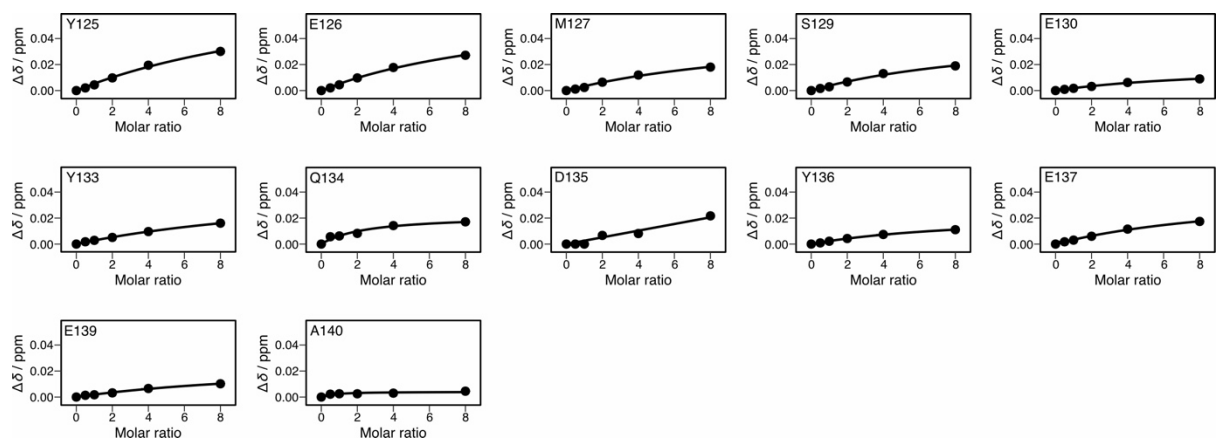

**Supplementary Fig. 3 | Titration curves of chemical shift changes for αSyn residues Y125–A140 upon addition of oxEGCG.** Points represent residue-specific weighted chemical shift differences ($\Delta\delta$), and solid lines indicate fits to a 1:1 binding model used to estimate apparent $K_D$ values. Error bars, which represent root-mean-square deviations (RMSDs), are smaller than the data-point symbols.

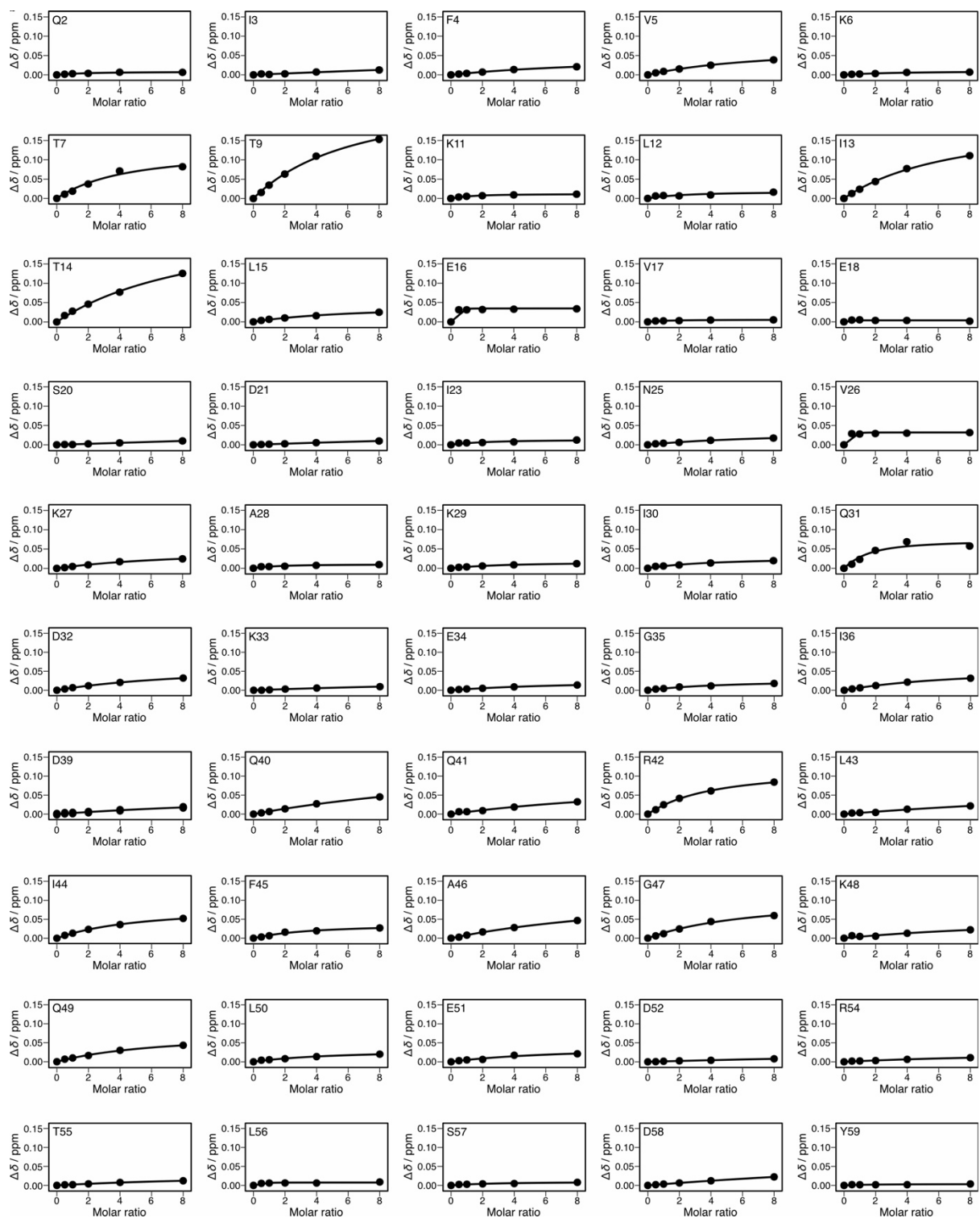

**Supplementary Fig. 4 | Titration curves of chemical shift changes for ubiquitin residues Q2–Y59 upon addition of oxEGCG.** Points represent residue-specific weighted chemical shift differences ($\Delta\delta$), and solid lines indicate fits to a 1:1 binding model used to estimate apparent $K_D$ values. Error bars, which represent root-mean-square deviations (RMSDs), are smaller than the data-point symbols.

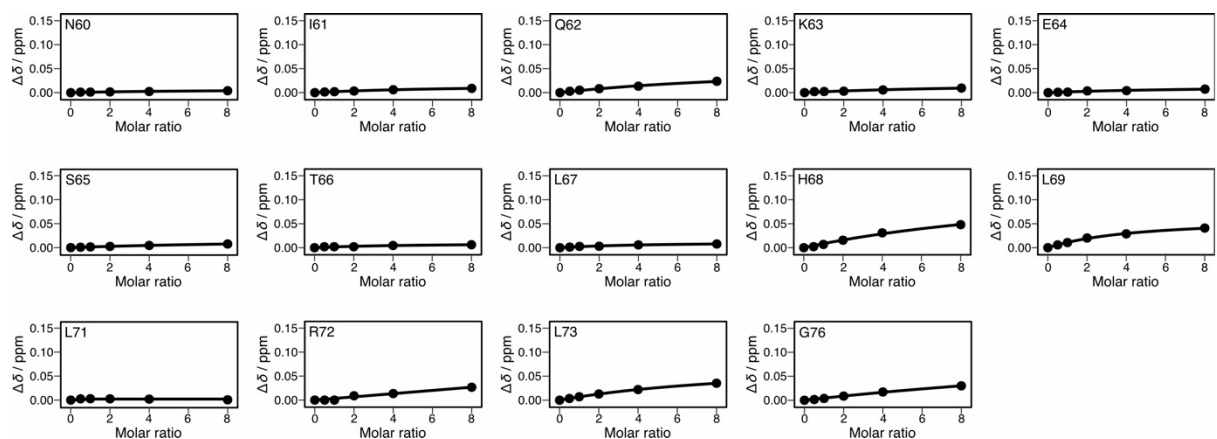

**Supplementary Fig. 5 | Titration curves of chemical shift changes for ubiquitin residues N60–G76 upon addition of oxEGCG.** Points represent residue-specific weighted chemical shift differences ($\Delta\delta$), and solid lines indicate fits to a 1:1 binding model used to estimate apparent $K_D$ values. Error bars, which represent root-mean-square deviations (RMSDs), are smaller than the data-point symbols.

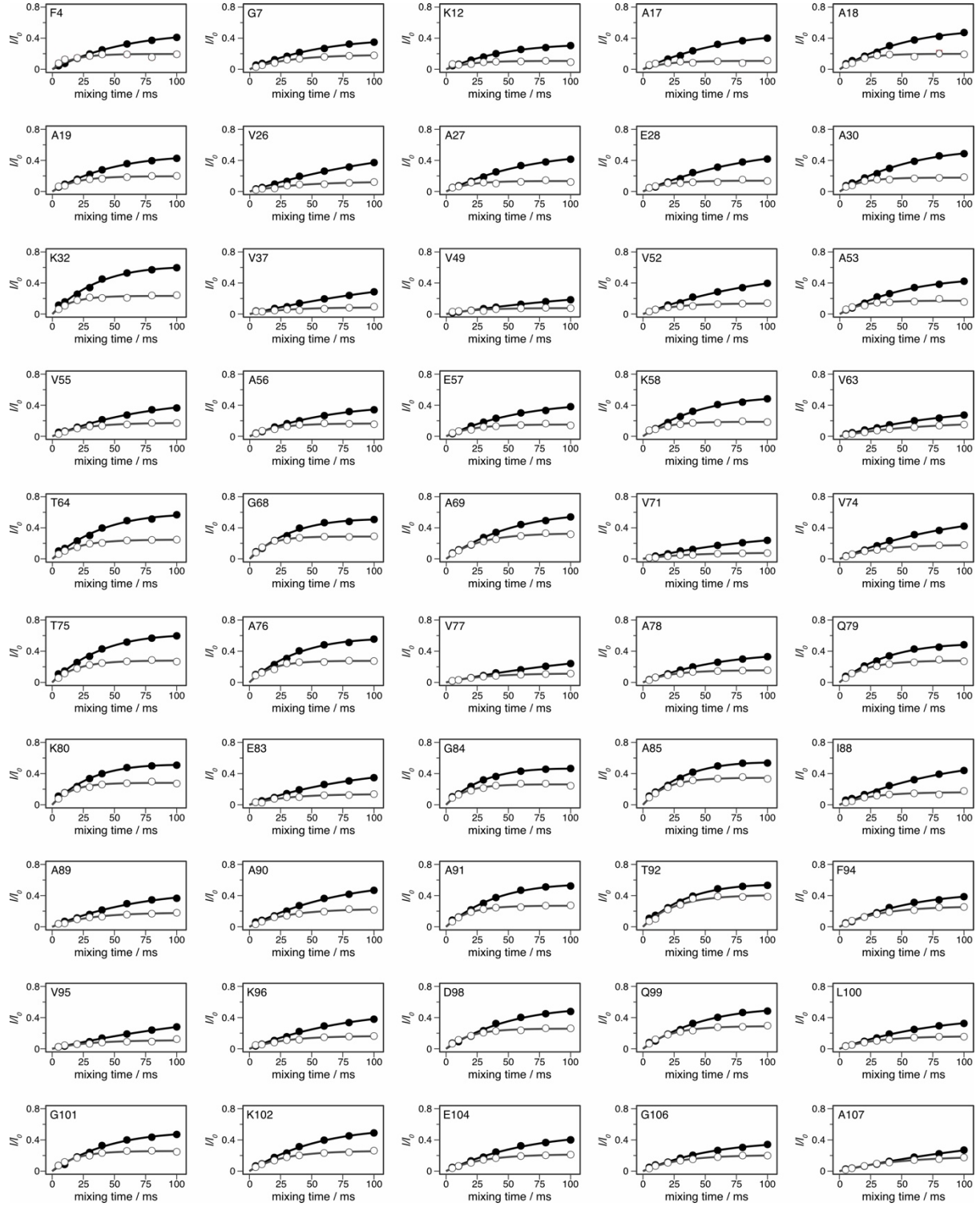

**Supplementary Fig. 6 | CLEANEX-PM buildup curves of αSyn residues F4–A107.** Normalized intensities are shown for αSyn in the absence (closed circles) and presence (open circles) of oxEGCG(80%). Solid lines indicate fits used to determine $k_{\text{IH}}$ values. Error bars, which represent root-mean-square deviations (RMSDs), are smaller than the data-point symbols.

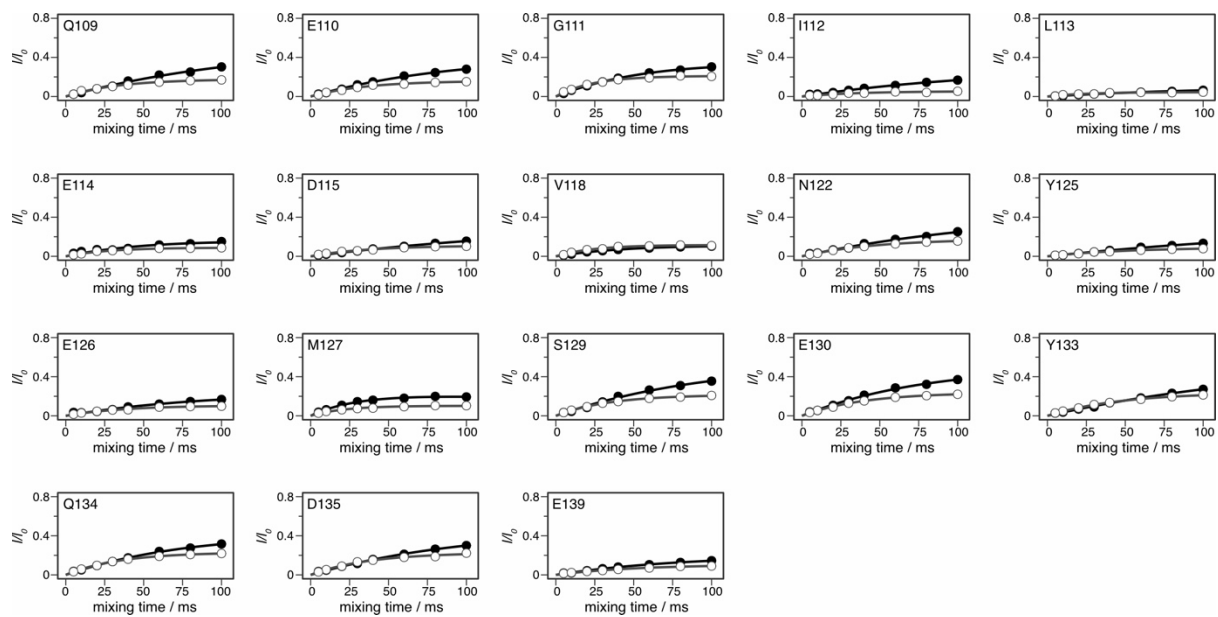

**Supplementary Fig. 7 | CLEANEX-PM buildup curves of αSyn residues Q109–E139.** Normalized intensities are shown for αSyn in the absence (closed circles) and presence (open circles) of oxEGCG(80%). Solid lines indicate fits used to determine $k_{1H}$ values. Error bars, which represent root-mean-square deviations (RMSDs), are smaller than the data-point symbols.

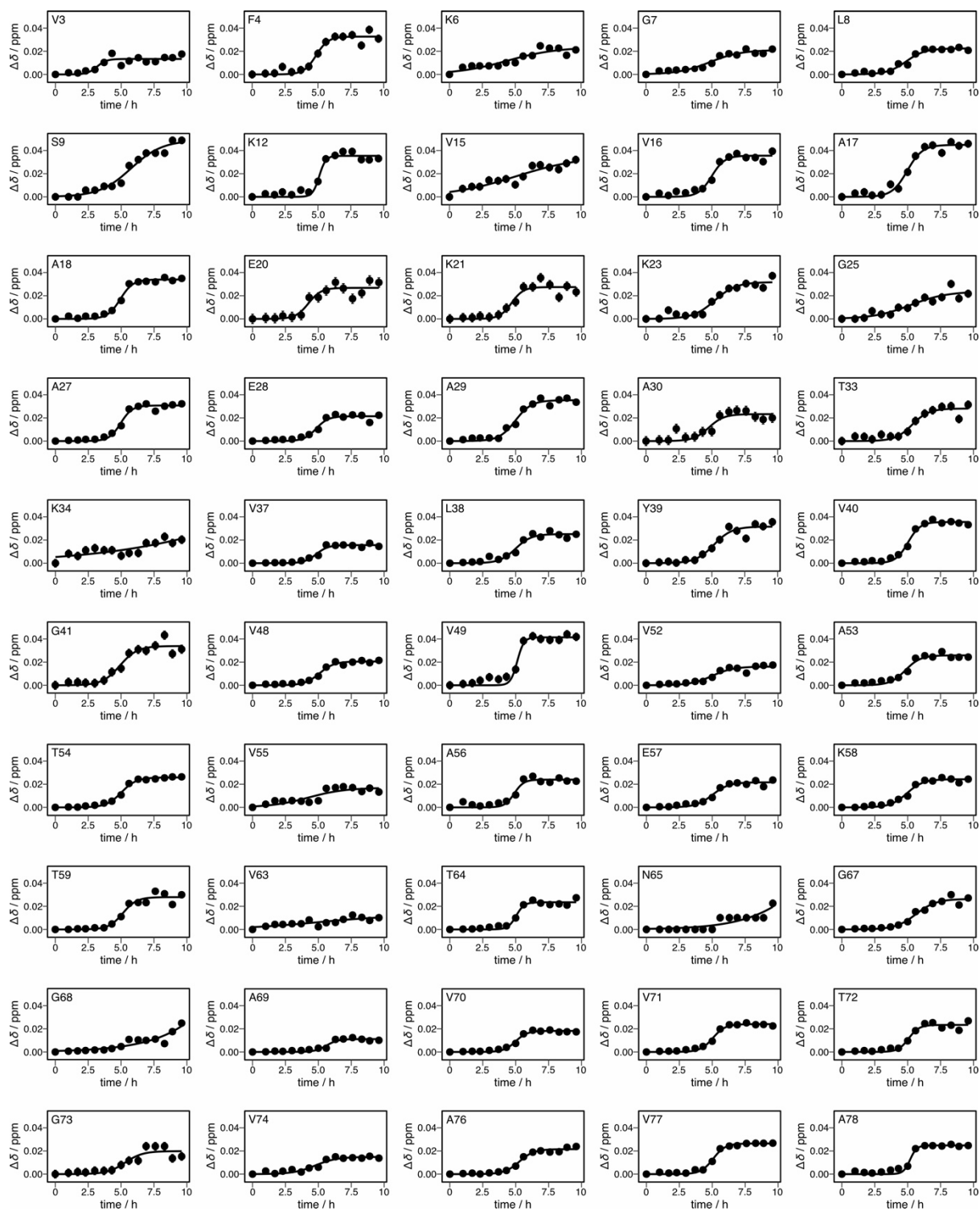

**Supplementary Fig. 8 | Residue-specific chemical shift changes of αSyn residues V3–A78 in the presence of oxEGCG(80%) under shear flow. Solid lines indicate sigmoidal fits used to determine $t_{1/2}$ values. Error bars indicate root-mean-square deviations (RMSDs).**

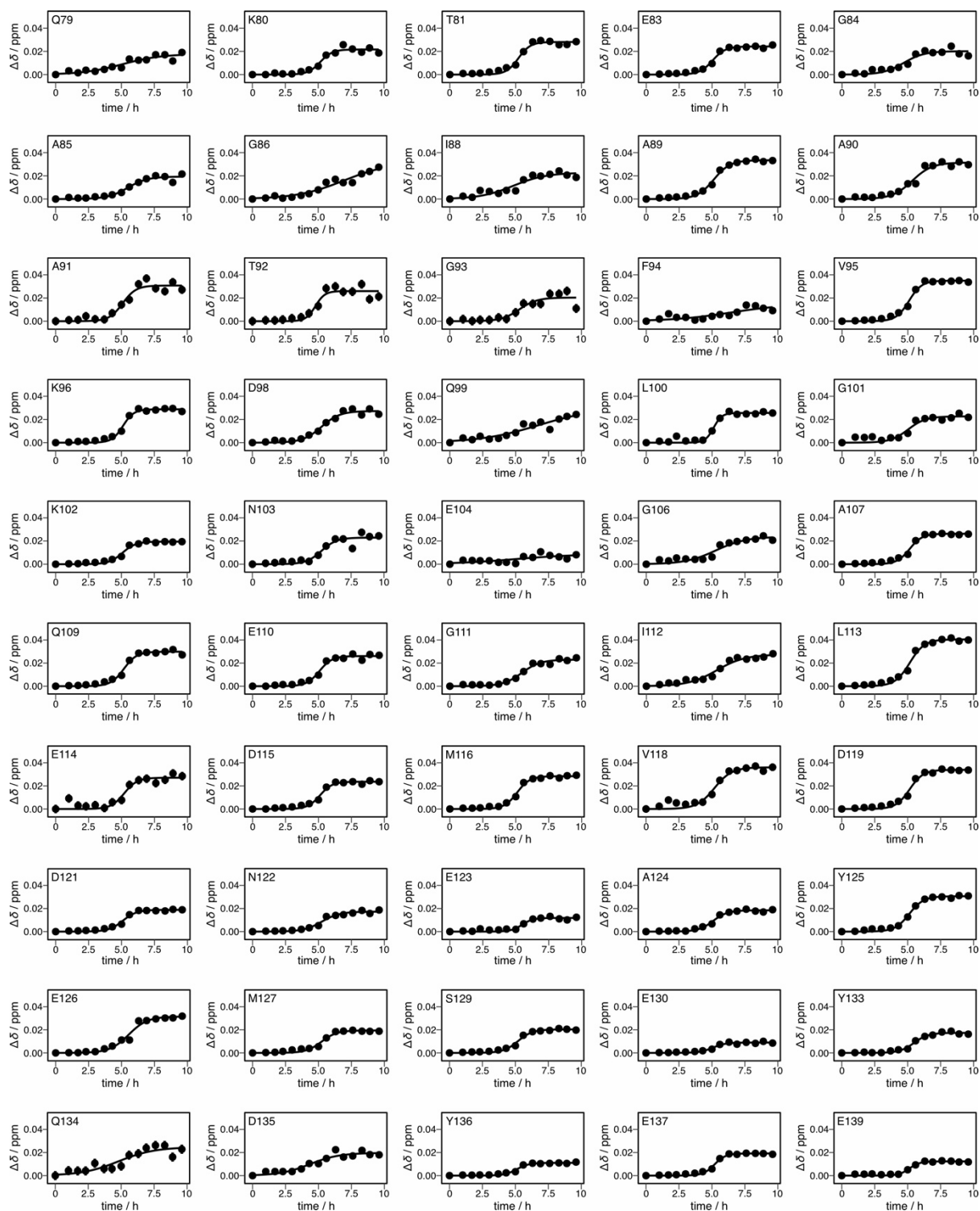

**Supplementary Fig. 9 | Residue-specific chemical shift changes of αSyn residues Q79–E139 in the presence of oxEGCG(80%) under shear flow.** Solid lines indicate sigmoidal fits used to determine $t_{1/2}$ values. Error bars, which represent root-mean-square deviations (RMSDs), are smaller than the data-point symbols.

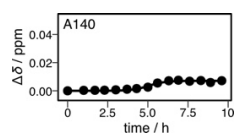

**Supplementary Fig. 10 | Chemical shift change of αSyn residue A140 in the presence of oxEGCG(80%) under shear flow.** Solid line indicates a sigmoidal fit used to determine $t_{1/2}$. Error bars, which represent root-mean-square deviation (RMSD), are smaller than the data-point symbols.

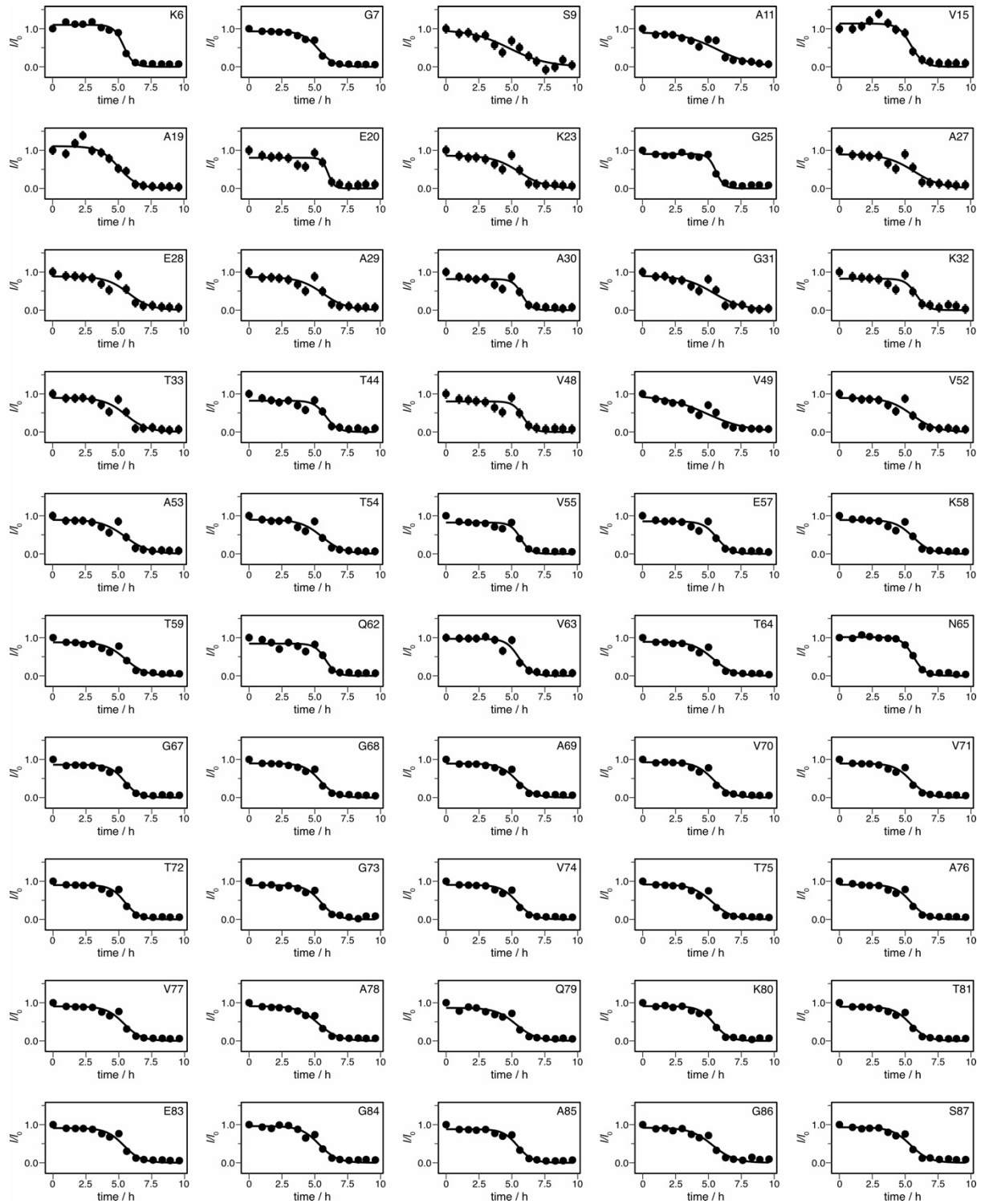

**Supplementary Fig. 11 | Residue-specific signal intensity decays of αSyn residues K6–S87 in the presence of oxEGCG(80%) under shear flow. Intensities are normalized to the initial value ($I/I_0$). Solid lines indicate sigmoidal fits used to determine $t_{1/2}$ values. Error bars indicate root-mean-square deviations (RMSDs).**

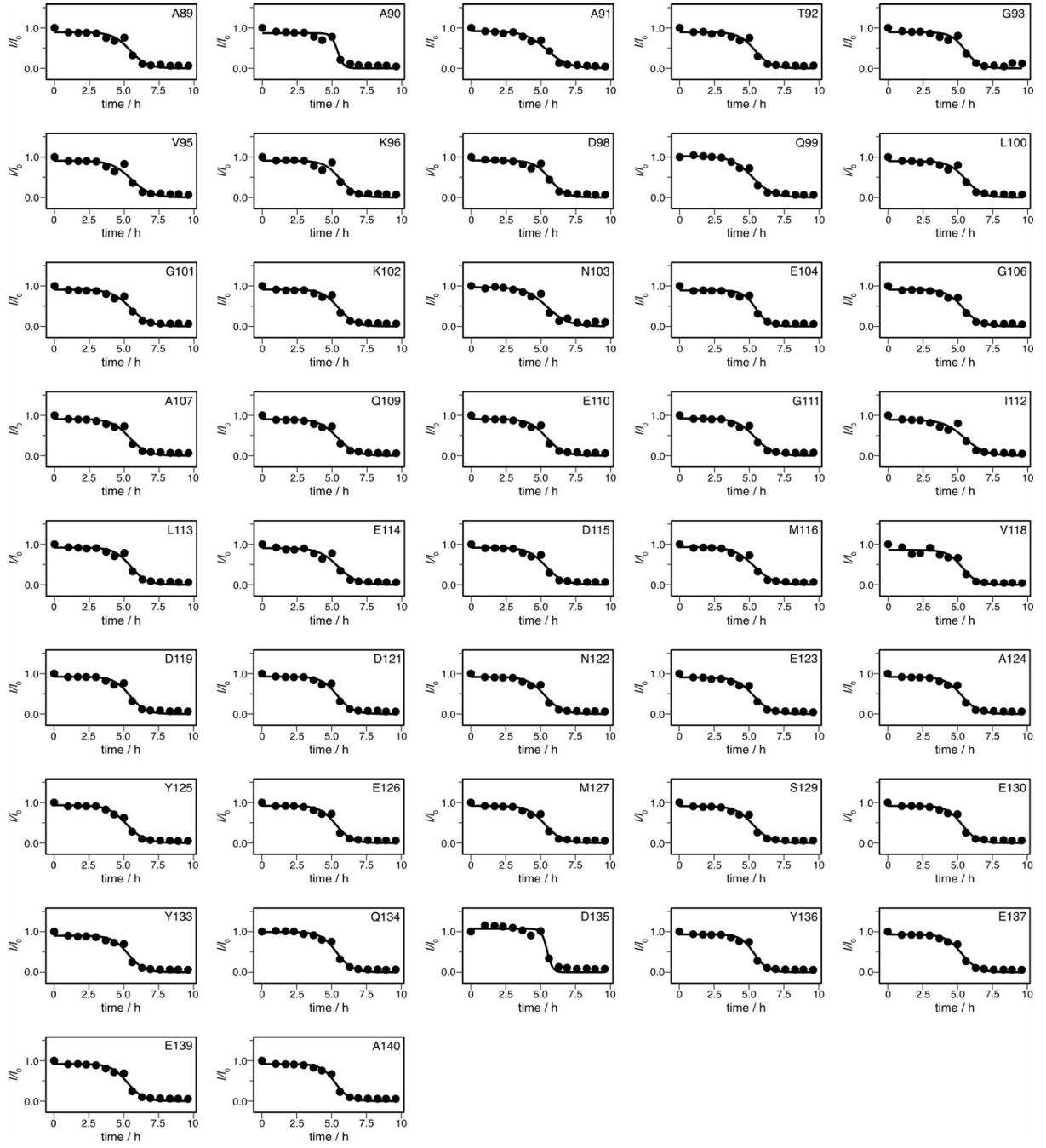

**Supplementary Fig. 12 | Residue-specific signal intensity decays of αSyn residues A89–A140 in the presence of oxEGCG(80%) under shear flow.** Intensities are normalized to the initial value ($I/I_0$). Solid lines indicate sigmoidal fits used to determine $t_{1/2}$ values. Error bars, which represent root-mean-square deviations (RMSDs), are smaller than the data-point symbols.
